# Plant pathogens cleave 2′cADPR to suppress TIR immune signaling

**DOI:** 10.64898/2026.06.15.732069

**Authors:** Ohad Roth, Aaron W. Lawson, Federica Locci, Talya Barak, Silvina Perin, Elke Logemann, Gil Amitai, Jane E. Parker, Paul Schulze-Lefert, Rotem Sorek

## Abstract

Small molecules produced by TIR (Toll-interleukin-1 receptor) domains are essential for plant immune signaling. A central TIR-derived signal is 2ʹcADPR, a cyclic ADP-ribose (ADPR) molecule that is generated by plant TIR-domain proteins upon sensing pathogen infection. Here we show that XopQ, a virulence protein widespread in plant pathogenic bacteria, cleaves 2ʹcADPR and converts it into an inactive, linear ADPR molecule. Steady-state kinetic analyses revealed that XopQ is a highly efficient enzyme that hydrolyzes 2ʹcADPR with near diffusion-limited catalytic efficiency, while displaying high specificity and no detectable activity toward closely related molecules such as 3ʹcADPR. We show that XopQ eliminates 2ʹcADPR *in planta* and prevents accumulation of pRib-AMP, a derivative of 2ʹcADPR that activates the downstream immune complex EDS1-PAD4. Our data demonstrate that XopQ inhibits PAD4-dependent plant immunity and can promote pathogen virulence even when the pathogen carries an avirulence effector that triggers TIR-mediated immune signaling. Finally, we show that XopQ can also inhibit type VI Thoeris, a bacterial anti-phage defense system that relies on 2ʹcADPR signaling, possibly explaining why XopQ and type VI Thoeris are not observed co-occurring in bacterial genomes. Our findings uncover a widespread and conserved strategy used by plant pathogens to directly target host TIR signaling.

## Introduction

Intracellular signaling through TIR-derived small molecules has recently emerged as a central aspect of the plant immune response^1^. In plants, the TIR (acronym of Toll-interleukin-1 receptor) domain is an enzyme whose main substrate is the molecule nicotinamide adenine dinucleotide (NAD^+^)^2,3^. When a plant TIR-domain protein detects the intracellular presence of a pathogen effector during pathogenesis, the TIR domain becomes active and generates a signaling molecule that typically contains ADP-ribose (ADPR)^1^. The molecule then activates a protein complex that includes the protein EDS1 to propagate the downstream immune response^1^.

Several TIR-derived immune signaling molecules have been characterized in plants in recent years^2–6^. One of the central signaling molecules, shown to be produced by both dicotyledonous and monocotyledonous TIRs, is 2ʹcADPR (also known as 1ʹʹ-2ʹ glycocyclic ADPR), in which ADPR is cyclized via a ribose-ribose bond^2,3^. It was hypothesized that 2ʹcADPR is further converted in the plant cell into a short-lived molecule called pRib-AMP, which then binds and activates a heterodimeric complex comprising the proteins EDS1 and PAD4^5,7^.

It is thought that TIR-mediated immune signaling first evolved in bacteria, where it serves to protect against bacteriophage (phage) infection^8^. TIR signaling is manifested in a family of bacterial defense systems collectively called Thoeris^8–12^. As in the case of plant TIRs, Thoeris TIR-containing proteins respond to phage infection by producing ADPR-based signaling molecules that activate a second protein in the Thoeris system to perform the downstream immune function^12^. Different members within the family of Thoeris defense systems produce distinct signaling molecules. These include cyclized forms of ADPR such as 3ʹcADPR^3,8^, N7-cADPR^10^ and canonical cADPR^12^, as well as ADPR linked to histidine (His-ADPR)^11^. A recent study described a bacterial Thoeris system that produces 2ʹcADPR as an immune signaling molecule, proposing a likely evolutionary origin for TIR-dependent immunity in plants^12^.

Recent studies showed that phages can overcome bacterial immunity by targeting TIR-derived signaling molecules^11,13–16^. Many phages express counter-defense proteins that bind and sequester the immune signaling molecules, allowing the phage to propagate even if the TIR-containing protein has detected its presence in the cell^13,16^. Phages were also shown to encode enzymes that cleave and inactivate bacterial immune signaling molecules produced by Thoeris and by other defense systems, including CBASS, and Pycsar^13,17–19^. Despite the centrality of TIR-mediated immunity in plants, it is currently unknown whether and how plant pathogens interfere with TIR signaling.

Here we report that XopQ, a virulence factor conserved and widespread across plant pathogens, is an enzyme that efficiently and specifically cleaves 2ʹcADPR. XopQ (*Xanthomonas* outer protein Q)^20^ is delivered into plant cells during infection by pathogenic bacteria from the *Xanthomonas*, *Pseudomonas*, *Acidovorax, Erwinia* and *Ralstonia* lineages, that cause numerous plant diseases including blight disease in rice, bacterial speck of tomato, bacterial fruit blotch in cucurbits, fire blight in apples, bacterial wilt in potato, and more^21,22^. XopQ was discovered more than two decades ago in functional screens aimed to detect virulence factors delivered by pathogens into plant cells and was predicted to be a nucleoside hydrolase based on sequence homology^20,23,24^. Since its discovery, XopQ has been shown to be essential for maximizing infection efficiency of plant pathogens such as *Xanthomonas oryzae* and *Pseudomonas syringae*^25–27^, but the mechanism by which XopQ confers a virulence advantage remained unclear. Our data show that XopQ efficiently breaks 2ʹcADPR into ADPR, thus inactivating the TIR-derived signaling molecule, preventing pRib-AMP accumulation, and suppressing EDS1-PAD4-mediated immunity. We show that XopQ is a highly efficient and specific 2ʹcADPR hydrolase, and does not target related molecules such as 3ʹcADPR. Our experiments demonstrate that XopQ cleaves 2ʹcADPR in vitro, in bacteria and *in planta* to inactivate TIR immune signaling, explaining a common and conserved counter-immunity mechanism employed by plant pathogens.

## Results

### Structural modeling predicts that XopQ specifically targets 2ʹcADPR

Sequence analysis suggested that XopQ is a member of a distinct family of nucleoside-hydrolyzing enzymes^20^, and mutations in the predicted active site of XopQ suppressed virulence^25,27^. However, early experiments with XopQ did not detect nucleoside or nucleotide hydrolase activity when this protein was incubated with either adenosine, guanosine, thymidine, cytidine, uridine, inosine, pseudouridine, GTP, UMP, or canonical cADPR^25,28,29^. Nevertheless, a crystal structure of XopQ from *Xanthomonas oryzae*, published in 2014 by Yu et al^28^, revealed a linear ADPR molecule bound in the predicted active site of XopQ. We initiated our study after noticing that the linear ADPR bound within the XopQ structure is bent in an angle that brings the two ribose moieties of ADPR in close proximity, generating a shape resembling that of 2ʹcADPR (Figure 1A). This led us to hypothesize that XopQ might specifically target 2ʹcADPR as a substrate.

**Figure 1:**
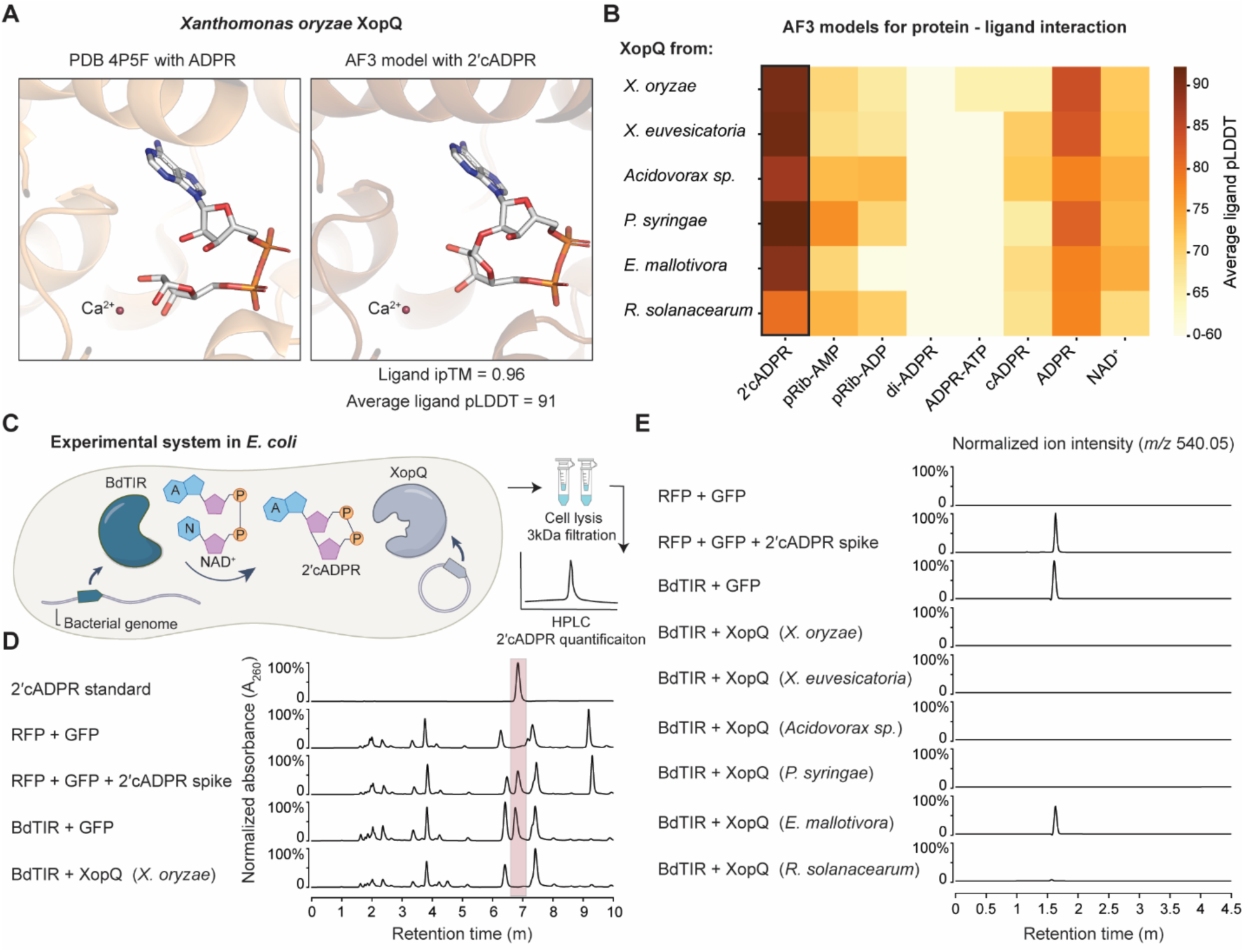
Bacterial XopQ targets the plant immune signaling molecule 2ʹcADPR. **A**. Comparison of the *X. oryzae* XopQ bound to ADPR (PDB 4P5F) and the Alphafold3-modeled XopQ bound to 2**ʹ**cADPR. Shown is a close-up view of the active site area. Presented ligand ipTM and pLDDT values were calculated as the average of 5 AlphaFold3 models. **B**. AlphaFold3-predicted ligand pLDDT values, averaged across ligand atoms, for the indicated molecules co-folded with XopQ homologs. **C**. Schematic illustration of the *E. coli*-based experimental system to assess XopQ activity toward BdTIR-generated 2**ʹ**cADPR. Bases (adenine, A, or nicotinamide, N) are in blue, ribose moieties in purple, and phosphate groups are in orange. Sample preparation included filtration of cell lysate to enrich for small molecules. **D.** HPLC absorbance chromatograms of filtered lysates from *E. coli* expressing the indicated proteins. Data are presented as absorbance (A_260_) values, with each chromatogram normalized independently to its maximum value. Representative of three or more replicates. **E.** HPLC-MS chromatograms of filtered lysates from *E. coli* expressing the indicated proteins. Data are presented as ion intensity for *m/z* 540.05, with each chromatogram normalized to the BdTIR + GFP control, except for the RFP + GFP + 2**ʹ**cADPR spike chromatogram that was normalized independently to its maximum value. Representative of three or more replicates.

We used AlphaFold3^30^ to co-fold 2ʹcADPR and XopQ. AlphaFold3 modeled 2ʹcADPR within the XopQ active site with high confidence (ipTM=0.96; average pLDDT of ligand atoms=0.91)(Figure 1A). The modeled 2ʹcADPR took a position and shape very similar to the linear ADPR detected within the solved XopQ crystal structure (Figure 1A; Figure S1). Co-folding XopQ with other known TIR-derived plant signaling molecules, including pRib-AMP, pRib-ADP, ADPR-ADPR (di-ADPR), and ADPR-ATP yielded substantially lower AlphaFold3 confidence scores for binding, suggesting specificity towards 2ʹcADPR (Figure 1B).

AlphaFold3 is known to be prone to false-positive predictions of protein-protein and protein-ligand interactions^31,32^. However, recent studies have shown that if the same interacting partner is consistently predicted with high confidence across homologs of a given protein, this increases chances that the predicted interaction represents real binding^32^. We therefore modeled 2ʹcADPR binding for XopQ homologs from diverse plant pathogens, including those of *Xanthomonas euvesicatoria*, *Pseudomonas syringae*, *Ralstonia solanacearum*, *Erwinia mallotivora* and *Acidovorax sp*. SUPP3334 (Table S1). Despite substantial sequence divergence between these homologs (Figure S2), AlphaFold3 consistently predicted high confidence binding of all XopQ homologs to 2ʹcADPR, with lower confidence scores predicted for interactions with other related molecules (Figure 1B). The 2ʹcADPR ligand was predicted to be bound in a nearly identical orientation within the putative binding sites of XopQ homologs, lending further support to the hypothesis that XopQ specifically binds 2ʹcADPR (Figure S3).

The structural modeling presented above raised the hypothesis that XopQ might sequester or cleave 2**ʹ**cADPR. To assess the likelihood of this hypothesis, we established an experimental system in *Escherichia coli* to test whether XopQ targets 2**ʹ**cADPR. Previous studies have shown that BdTIR, a TIR-domain protein from the monocotyledonous plant *Brachypodium distachyon*, produces substantial amounts of 2**ʹ**cADPR when expressed in *E. coli*^2,15^. Our experiments reproduced this observation, showing that 2**ʹ**cADPR could readily be detected in lysates of *E. coli* cells expressing BdTIR (Figures 1C, 1D), and liquid chromatography followed by mass spectrometry (LC-MS) further verified the identity of the molecule (Figure 1E). However, when BdTIR was expressed with XopQ, 2**ʹ**cADPR could not be detected in the cell lysates (Figure 1D, 1E). Homologs of XopQ from *X. euvesicatoria*, *P. syringae*, *Acidovorax sp* and *R. solanacearum* also led to complete or near-complete elimination of 2**ʹ**cADPR from BdTIR-expressing cells, while the *E. mallotivora* homolog caused only partial reduction of 2**ʹ**cADPR accumulation (Figure 1E). Together, these results suggest a conserved 2**ʹ**cADPR targeting capabilities among XopQ proteins from diverse plant pathogens.

### XopQ catalyzes 2ʹcADPR hydrolysis

XopQ belongs to a family of enzymes annotated as “inosine-uridine preferring nucleoside hydrolases” (Pfam accession: PF01156). These calcium-dependent enzymes catalyze the hydrolysis of purine and pyrimidine nucleosides, separating the base from the associated ribose^33^. To test whether XopQ catalyzes 2ʹcADPR in a similar manner, we expressed three XopQ homologs and purified them for biochemical characterization (Table S1). When incubated with 2ʹcADPR at room temperature, all three purified XopQ proteins rapidly and completely consumed the molecule (Figure 2A). Our data show that XopQ converts 2ʹcADPR into a single product with mass and retention time matching that of ADPR, confirming that XopQ is a 2ʹcADPR cleaving enzyme (Figures 2A, 2B). Point mutations in two conserved aspartic acid residues predicted to orient 2ʹcADPR and Ca^2+^ in the active site abolished enzymatic activity (Figure 2A; Figure S2).

**Figure 2.**
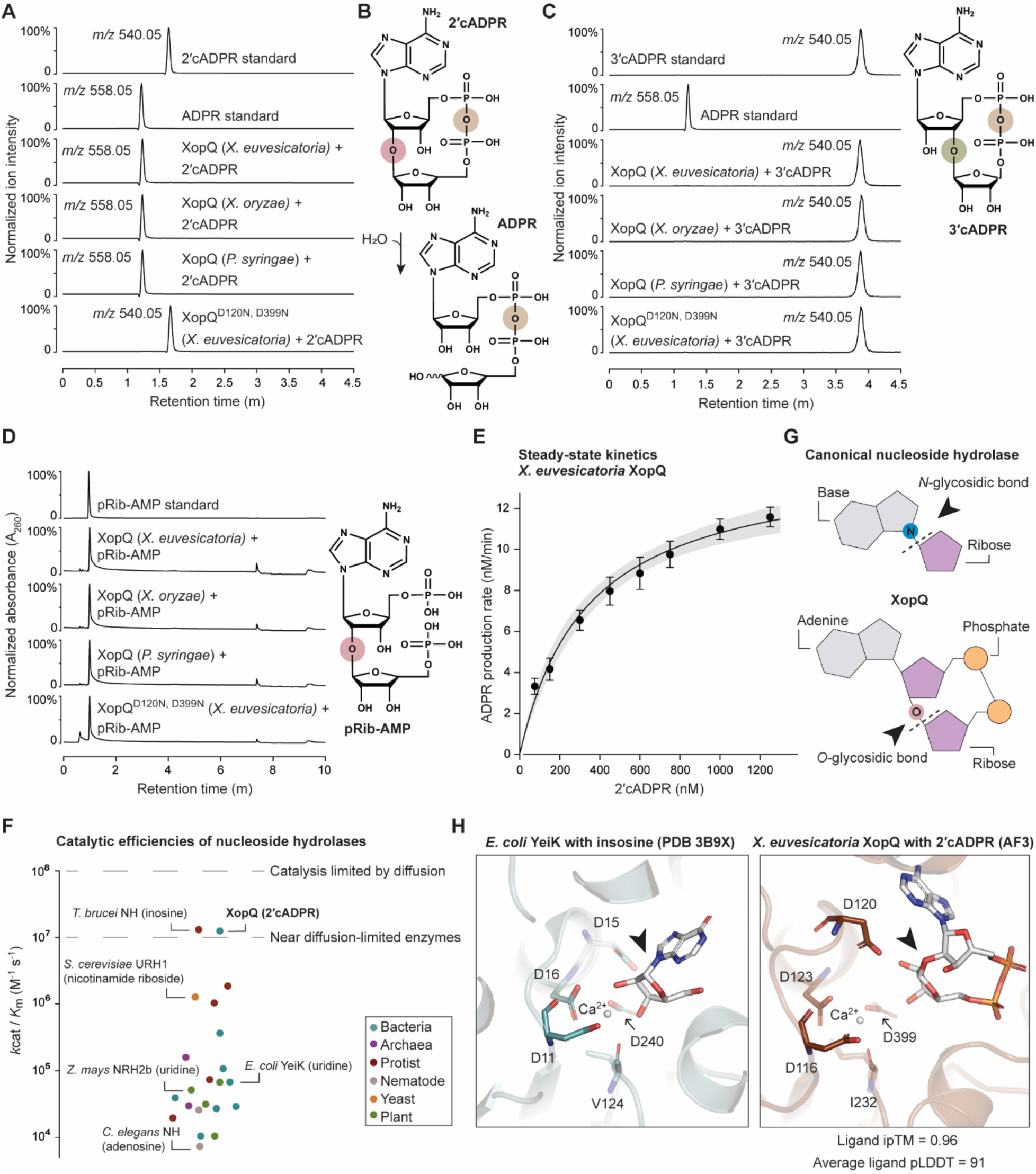
XopQ hydrolyzes 2ʹcADPR to ADPR with high efficiency and specificity. **A**. HPLC-MS chromatograms of in vitro reactions of purified XopQ proteins from the indicated bacterial species, incubated with 2ʹcADPR for 20 minutes at 25 °C. Reactions contained 100 nM protein and 10 µM 2ʹcADPR. Data are shown as ion intensity for *m/z* 540.05 (2ʹcADPR) and *m/z* 558.05 (ADPR), with each chromatogram normalized independently to its maximum value. Presented chromatograms are representatives of at least three replicates. XopQ proteins were cloned without the N-terminal signal peptide marking the protein for secretion (Table S1). **B.** Schematic representation of the reaction catalyzed by XopQ. The hydrolyzed 1ʹʹ-2ʹ *O*-glycosidic bond is in pink and the pyrophosphate bond is in brown. **C, D.** HPLC chromatograms of in vitro reactions of purified XopQ proteins from the indicated bacterial species, incubated with 3ʹcADPR (**C**) or pRib-AMP (**D**) for 20 minutes at 25 °C. Reactions contained 100 nM protein and 10 µM of the indicated substrate. Data are presented as normalized ion intensity for *m/z* 540.05 and *m/z* 558.05 (**C**), or normalized absorbance at A_260_ (**D**). Each chromatogram was normalized independently to its maximum value. Presented chromatograms are representatives of at least three replicates. **E.** Steady-state kinetics of 2ʹcADPR-to-ADPR hydrolysis by *X. euvesicatoria* XopQ. Reactions contained ∼0.05 nM protein and were performed at 25 °C. Each point represents the mean of three independent replicates; error bars represent standard error of the mean. Shaded region indicates the 95% confidence interval of the fitted curve. **F.** Comparison of the catalytic efficiency parameter (*k*_cat_/*K*_m_) calculated here for XopQ and previously characterized nucleoside hydrolases. Parameters of previously studied enzymes with their preferred substrates and conditions were retrieved from the BRENDA database^35^. **G.** Schematic illustrations comparing the reactions catalyzed by canonical nucleoside hydrolases and XopQ. Arrowheads indicate the cleaved bonds. **H.** Structural comparison of the pyrimidine nucleoside hydrolase YeiK from *E. coli* bound to inosine^36^ (PDB 3B9X) and *X. euvesicatoria* XopQ bound to 2ʹcADPR (AlphaFold3 model). Conserved residues that coordinate Ca^2+^ and position the substrates in the active site are shown as sticks. Arrowheads indicate the cleaved bonds.

Notably, two of the XopQ proteins we purified and tested – from *X. oryzae* and *P. syringae* (the latter is also known as HopQ1) – have previously been reported to be catalytically inactive when tested in vitro against a panel of nucleosides and nucleotides^25,29^. This further suggests that XopQ specifically prefers 2ʹcADPR as substrate. To examine the specificity of XopQ in more detail, we tested its ability to target related molecules. We found that none of the three tested XopQ homologs could hydrolyze 3ʹcADPR (Figure 2C), a molecule very similar to 2ʹcADPR which differs only in the location of the *O-*glycosidic bond connecting the two ribose moieties. We observed a rapid and complete 2ʹcADPR to ADPR hydrolysis by XopQ even in the presence of a 1000-fold excess of 3ʹcADPR over 2ʹcADPR, suggesting that 3ʹcADPR does not compete with 2ʹcADPR for binding to XopQ (Figure S4).

The plant immune signal pRib-AMP is another molecule whose structure and composition closely resemble that of 2’cADPR (Figure 2D). pRib-AMP and 2ʹcADPR share the same *O*-glycosidic bond that connects the two ribose moieties via carbon C2ʹ, but in 2ʹcADPR the two phosphate groups are linked, while in pRib-AMP they are separated (Figure 2D). Despite the similarity between the two molecules and the presence of the same *O*-glycosidic bond, none of the XopQ homologs we tested cleaved pRib-AMP (Figure 2D). In our hands, XopQ was also incapable of cleaving the cyclic nucleotides 2ʹ3ʹcAMP and 2ʹ3ʹcGMP even after overnight incubation, despite earlier reports suggesting these molecules as XopQ substrates^34^ (Figure S5). In addition, XopQ did not cleave canonical cADPR, as previously reported^28^ (Figure S5C). Together, these results indicate that XopQ enzymes have evolved to selectively bind and hydrolyze the immune molecule 2ʹcADPR.

To assess the kinetic properties of XopQ-catalyzed 2ʹcADPR hydrolysis, we measured the initial rates of ADPR formation by XopQ across substrate concentrations ranging from 75 nM to 12.5 µM (Figure 2E). Steady-state kinetic analysis revealed that XopQ hydrolyzes 2ʹcADPR with near diffusion-limited catalytic efficiency (*k*_cat_/*K*_m_ = 1.43 X 10^7^ M^-1^ s^-1^) associated with a sub-micromolar Michaelis constant (*K*_m_ = 336 nM, 95% confidence interval 231-441 nM) (Figure 2E). XopQ showed a higher catalytic efficiency as compared to other nucleoside hydrolases from the PF01156 protein family (Figure 2F). Comparison of steady-state parameters for biochemically characterized enzymes of this family from animals, plants, protists, fungi and prokaryotes, showed that nearly all of them, when acting on their preferred nucleoside substrates, have catalytic efficiencies (*k*_cat_/*K*_m_) that are 10-1000 times lower than that of XopQ (Figure 2E, Table S2). These enzymatic properties of XopQ fit the hypothesis that its role as a virulence factor is to efficiently bind and eliminate a host immune-related signal present at low intracellular concentrations.

Notably, while all previously characterized nucleoside hydrolases from the PF01156 family cleave *N*-glycosidic bonds to remove a base covalently attached to a ribose moiety, XopQ catalysis cleaves the *O*-glycosidic bond of 2ʹcADPR, separating the ribose-ribose bond and linearizing the molecule into ADPR (Figure 2G). Comparison of the experimentally resolved structure of canonical nucleoside hydrolases to the AlphaFold3-predicted model of XopQ bound to 2ʹcADPR revealed a similar ligand orientation, in which the *O*-glycosidic bond of 2ʹcADPR occupies a position analogous to the *N*-glycosidic bond of canonical hydrolase substrates (Figure 2H). Together, our results establish XopQ virulence proteins as a new class of *O*-glycosyl hydrolases that likely evolved from *N*-glycosyl hydrolases by repurposing the active site to efficiently and specifically degrade the plant TIR-derived immune signal 2ʹcADPR.

### XopQ suppresses 2ʹcADPR-dependent immunity in bacteria

We recently identified a bacterial anti-phage system that utilizes 2ʹcADPR as an immune signaling molecule^12^. This system, denoted type VI Thoeris, is composed of a TIR-domain protein and a membrane-spanning protein called ThsG. It was demonstrated that upon phage infection, the Thoeris TIR protein produces 2ʹcADPR, which activates ThsG to induce bacterial growth arrest^12^. As shown for other Thoeris systems, induction of bacterial growth arrest prevents the phage from completing its replication cycle and limits the spread of infection to neighboring cells^8,10,11^.

To test whether XopQ can inhibit 2ʹcADPR-mediated immunity, we introduced plasmids expressing either wild-type XopQ or a catalytically inactive mutant (XopQ^D120N^) into an *E. coli* strain harboring this defense system (Figure 3A, Figure S6). We then assessed the ability of the Thoeris system to defend against the phage SECphi18 (Figure 3A). As previously shown^12^, phage SECphi18 formed 100-1000 fold fewer plaques on cells expressing the Thoeris system, confirming the substantial defense conferred by the system (Figure 3B). However, cells expressing both Thoeris and XopQ were not protected from SECphi18, and phages infecting these cells formed almost the same number of plaques as formed on a bacterial strain lacking the defense system (Figure 3B). Conversely, bacteria co-expressing Thoeris and the XopQ^D120N^ mutant maintained full resistance to phage infection, indicating that the catalytic activity of XopQ is required to inhibit Thoeris defense (Figure 3B).

**Figure 3.**
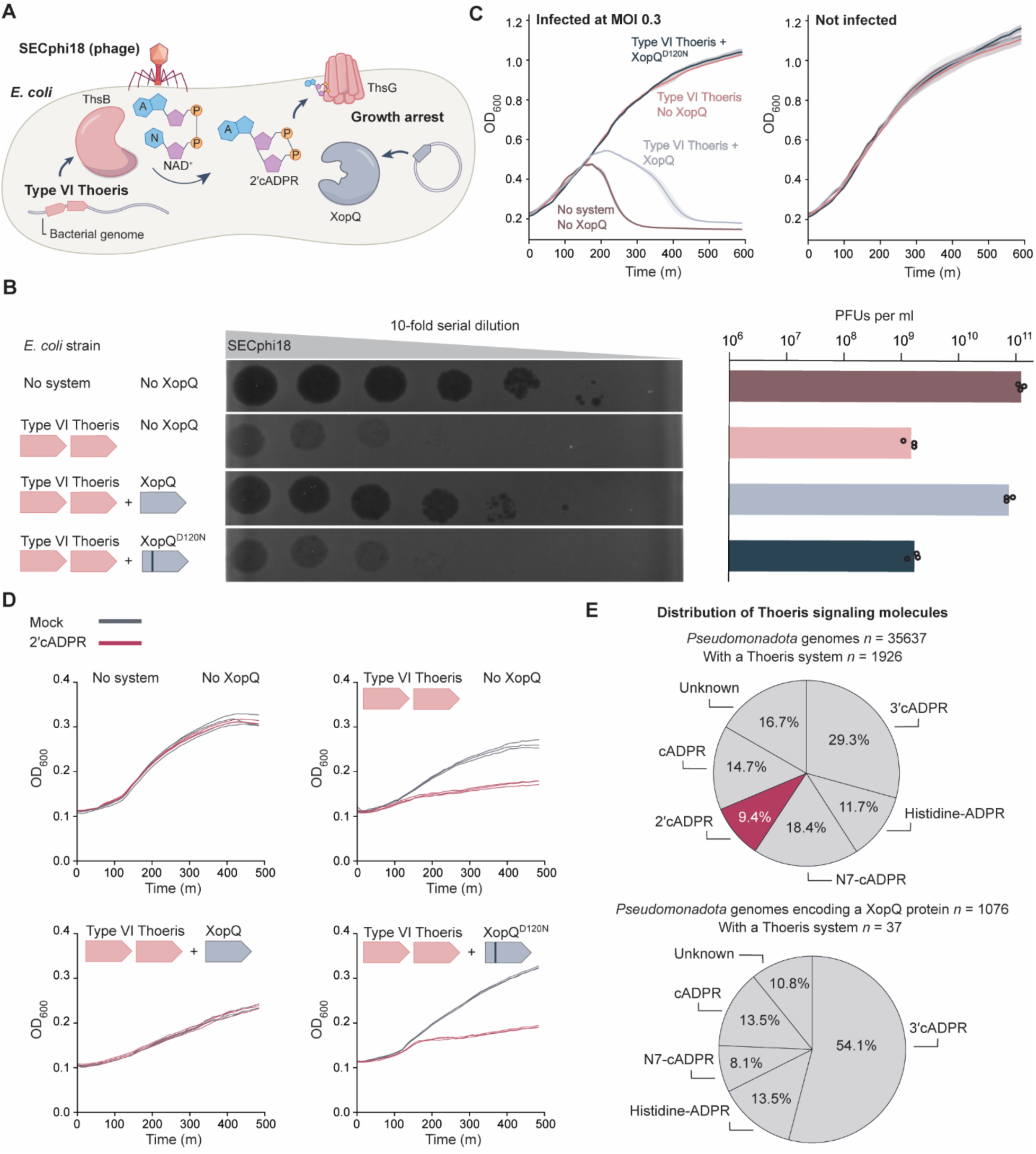
XopQ suppresses 2ʹcADPR-dependent immunity in bacteria. **A.** Schematic illustration of the experimental system used to assess the ability of XopQ to suppress anti-phage defense by the type VI Thoeris defense system in *E. coli*. **B.** Plaque assays assessing the efficiency of infection of phage SECphi18 on bacteria growing on a petri dish. Shown are 10-fold serial dilution plaque assays, using phage titers ranging from ∼10^6^ to ∼10^11^ plaque-forming units (PFUs) per ml. Left, representative images of plaque formation; right, quantification of PFUs per ml, showing average of three independent replicates with individual data points overlaid. **C.** Infection outcome in liquid media. Indicated strains were infected by phage SECphi18 at multiplicity of infection (MOI) of 0.3 (left) or grown in the absence of phage (right) at 25 °C. Each curve represents the mean of three independent experiments. Curve shading represents the standard error of the mean. **D**. XopQ suppresses 2ʹcADPR toxicity in cells expressing type VI Thoeris. Growth curves of *E. coli* cells expressing the type VI Thoeris system, with or without co-expressed XopQ variants. Media was supplemented with 1 mM 2ʹcADPR or H_2_O (Mock). Three repeats for each strain are shown as individual curves. **E.** Distribution of Thoeris systems categorized by their associated signaling molecules across *Pseudomonadota* genomes. Fisher’s exact test was used to assess whether Type VI Thoeris systems are significantly underrepresented in XopQ-encoding genomes (*P =* 0.03).

Experiments in which bacterial cultures were infected in liquid media confirmed the plaque assay results. In these experiments, bacteria were infected by phages at low multiplicity of infection (MOI) and then grown in liquid media. While cultures of Thoeris-expressing cells demonstrated growth even when infected by the phage, cells co-expressing Thoeris and XopQ, but not XopQ^D120N^, were lysed by the phage following infection (Figure 3C). These results further substantiate that XopQ inhibits 2ʹcADPR-mediated immunity.

To confirm that XopQ interferes with immune signaling downstream to the production of 2ʹcADPR, we applied exogenous 2ʹcADPR to cells expressing Thoeris. As shown previously, exogenous 2ʹcADPR can enter bacterial cells and cause the Thoeris ThsG protein to inhibit bacterial growth even in the absence of phage infection^12^ (Figure 3D). However, 2ʹcADPR did not affect growth in Thoeris-expressing cells that also expressed XopQ, suggesting that XopQ eliminated 2ʹcADPR within the cell (Figure 3D). Here, too, this activity depended on an intact active site, as the catalytically inactive mutant XopQ^D120N^ failed to curb 2ʹcADPR-mediated toxicity (Figure 3D). Together, these results demonstrate that XopQ can suppress TIR-mediated immunity through elimination of the signaling molecule 2ʹcADPR.

Thoeris systems in bacteria were shown to produce a variety of signaling molecules, with systems generating 2ʹcADPR accounting for roughly 10% of Thoeris systems in nature^12^. Our results suggest that XopQ might interfere with the capacity of 2ʹcADPR-dependent Thoeris to defend against phages in native conditions. In support of this hypothesis, none of the *Pseudomonadota* genomes in which we found a XopQ homolog (*n* = 1,076) also encoded a type VI Thoeris (Figure 3E, Table S3). Thoeris systems employing all other known signaling molecules (3ʹcADPR, cADPR, N7-cADPR, His-ADPR) did co-occur with XopQ in *Pseudomonadota* genomes, suggesting a specific depletion of 2ʹcADPR-utilizing systems from XopQ-encoding genomes.

### XopQ inactivates 2ʹcADPR *in planta*

Our data show that XopQ efficiently cleaves 2ʹcADPR both in vitro and in bacteria. Since the functional environment of XopQ as a virulence factor is within plant cells, we set out to examine whether XopQ can eliminate 2ʹcADPR when expressed *in planta*. For this, we induced the production of 2ʹcADPR in leaves of *Nicotiana benthamiana* using *Agrobacterium*-mediated transient expression of BdTIR (Figure 4A). In this experiment we relied on the previous observation that BdTIR constitutively produces 2ʹcADPR when expressed in *N. benthamiana*^2,37^. To prevent activation of *N. benthamiana* immunity and the resulting cell death triggered by BdTIR^2^, this experiment was performed in an immune-deficient *N. benthamiana* line lacking EDS1, PAD4, SAG101A, and SAG101B^38^. In addition to expressing BdTIR, we introduced WT XopQ or a catalytically inactive XopQ variant into the same *N. benthamiana* leaves and quantified 2ʹcADPR in leaf extracts 48 hours post *Agrobacterium* infiltration (Figure 4A; Figures S7).

**Figure 4.**
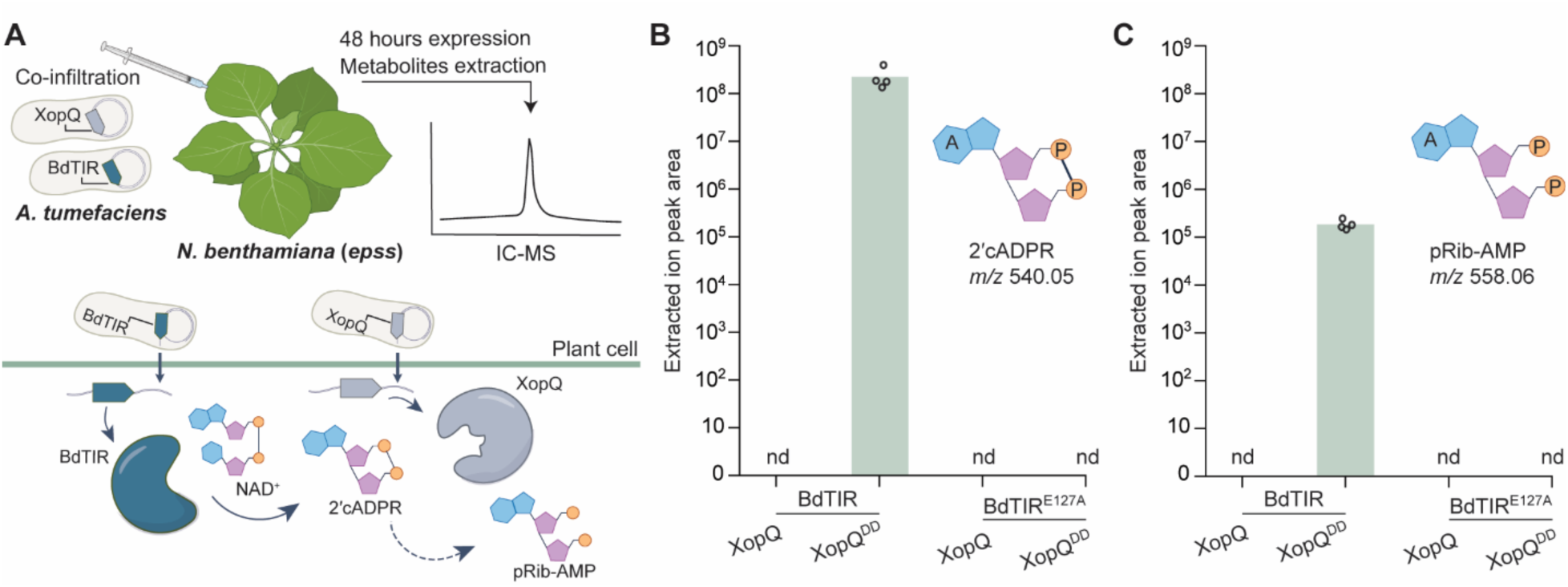
XopQ-mediated depletion of 2ʹcADPR and pRib-AMP *in planta*. **A.** Schematic illustration of the experimental system used to assess XopQ activity *in planta*. *A. tumefaciens* cultures carrying BdTIR or XopQ (WT or mutant) were mixed co-infiltrated into leaves of *N. benthamiana* deficient in EDS1, PAD4, SAG101A, and SAG101B (a quadruple mutant also denoted *epss*). Leaves were harvested for metabolite extraction 48 hours post infiltration. Dashed arrow indicates the hypothetical conversion of 2ʹcADPR to pRib-AMP. **B-C.** Extracted ion peak areas of 2ʹcADPR (*m/z* 540.05, panel **B**) and pRib-AMP (*m/z* 558.06, panel **C**) in samples obtained from leaves expressing the indicated BdTIR and XopQ constructs. XopQ^DD^ refers to the catalytically inactive XopQ^D120N, D399N^. Data are presented as the mean of four independent replicates, with individual data points overlaid; nd, not detected.

As expected, co-expression of BdTIR together with a catalytically inactive mutant of XopQ resulted in substantial accumulation of 2ʹcADPR in leaf extracts (Figure 4B). Experiments in which a catalytically inactive BdTIR was co-expressed with XopQ confirmed that 2ʹcADPR accumulation depended on BdTIR (Figure 4B). Remarkably, when WT XopQ was co-expressed with BdTIR, no 2ʹcADPR was detected in leaf extracts (Figure 4B). These results confirm that XopQ is enzymatically active *in planta* and can eliminate 2ʹcADPR from plant cells.

It was previously hypothesized that 2ʹcADPR can be converted in plant cells to pRib-AMP, which then triggers the immune response^7^. To date, pRib-AMP has not been observed in its free form in plant extracts and has only been detected bound to the EDS1–PAD4 protein complex, which presumably protects it from degradation^5,7,39^. After developing a chromatography protocol that enables distinguishing between pRib-AMP and its isomer ADPR (Figure S8), we were able to detect a molecule with mass and retention time matching the pRib-AMP standard in extracts obtained from leaves expressing BdTIR together with catalytically inactive XopQ (Figure 4C; Supplementary dataset S1). pRib-AMP could not be detected in leaf extracts when BdTIR was co-expressed with WT XopQ (Figure 4C). Together with our earlier observation that XopQ is specific for 2ʹcADPR and does not cleave pRib-AMP in vitro, these results support earlier findings^7^ by providing further evidence that pRib-AMP is generated downstream of 2ʹcADPR following TIR activation in plant cells, and establish XopQ as a suppressor of this immune pathway.

### Enhancement of *P. syringae* virulence via suppression of TIR-mediated immunity

We next sought to examine whether the ability of XopQ to degrade 2ʹcADPR confers plant pathogens with a virulence advantage. For this, we used the plant pathogen *P. syringae*, which encodes the XopQ homolog HopQ1, shown above to cleave 2ʹcADPR (Figure 2A). Certain strains of *P. syringae* also encode the effector protein AvrRps4, which is recognized in *Arabidopsis thaliana* Col-0 by the TIR-containing proteins pair RRS1-RPS4^40–42^, triggering the enzymatic activity of the RPS4 TIR domain^43^.

To ask whether HopQ1 can suppress AvrRps4-triggered immunity in *A. thaliana*, we generated operons expressing AvrRps4 together with WT or catalytically inactive HopQ1, and introduced them into a *P. syringae* pv. tomato DC3000 strain mutated in *hopQ1* and naturally lacking *avrRps4* (Figure 5A, 5B; Table S4)^44^. We then performed infection experiments in which these strains were spray-inoculated onto leaves of *A. thaliana* Col-0 and measured bacterial proliferation in these leaves four days after *P. syringae* inoculation. As expected, bacterial strains carrying operons expressing AvrRps4 proliferated ∼50-fold less efficiently compared to those carrying control RFP, in agreement with previous observations that AvrRps4 promotes RPS4-mediated immunity^40–42^ (Figure 5C; Figure S9A; Table S5). However, no difference in bacterial growth was observed for the strain carrying WT HopQ1 compared with the respective strain carrying catalytically inactive HopQ1 (Figure 5C; Figure S9A; Table S5). These results indicate that the presence of HopQ1 is insufficient to confer AvrRps4-expressing bacteria with a virulence advantage during colonization of WT Col-0 *A. thaliana* plants.

**Figure 5.**
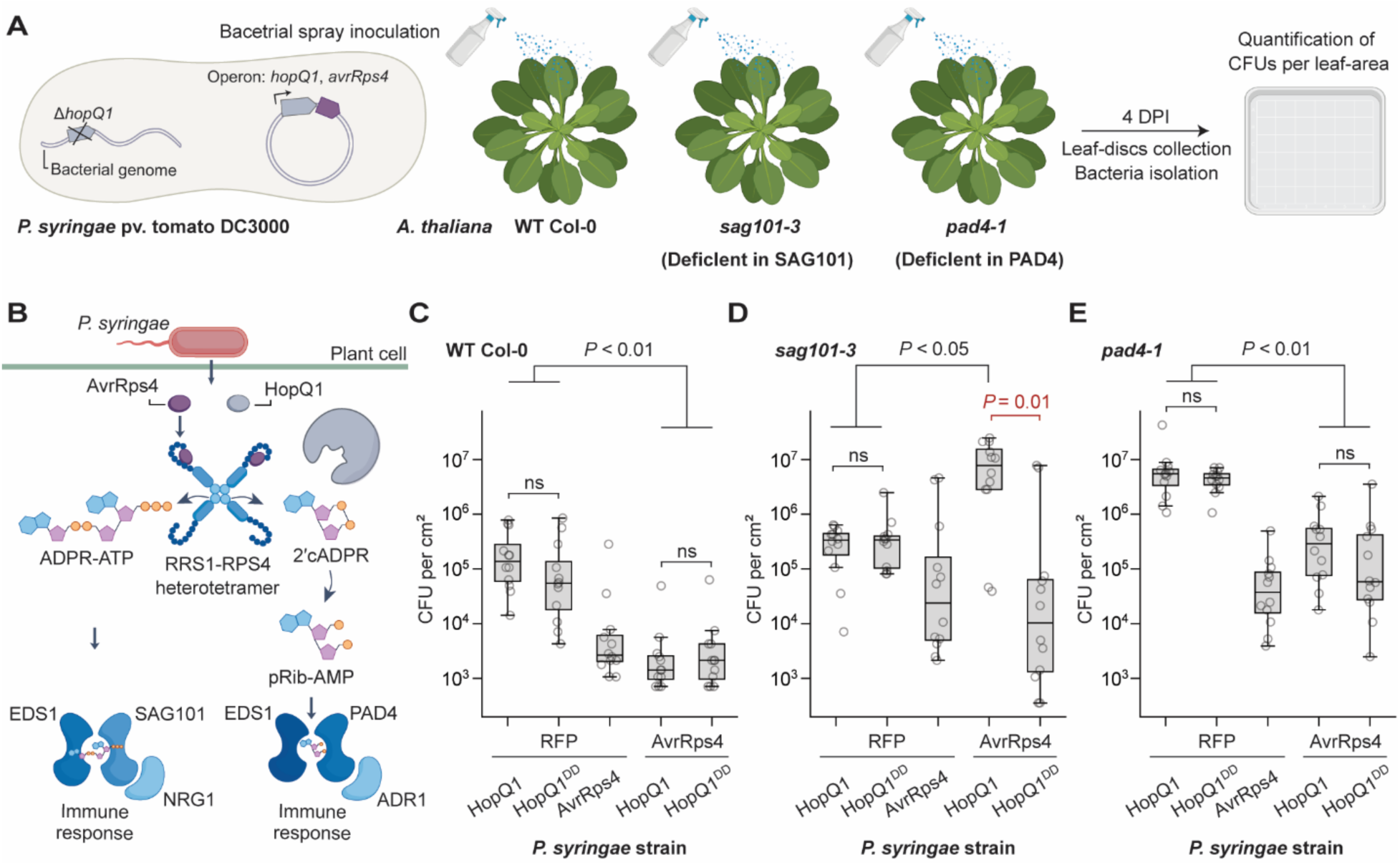
Pathogen-delivered HopQ1 suppresses TIR-mediated, PAD4-dependent immunity in *A. thaliana.* **A.** Schematic illustration of the experimental system used to assess the contribution of HopQ1 catalytic activity to suppression of AvrRps4-triggered immunity in *A. thaliana*. Operons contained native regulatory elements of *hopQ1* (promoter and terminator) and *avrRps4* (ribosome-binding site). **B**. An illustration showing that the same TIR-domain protein can produce distinct molecules that activate two immune nodes. AvrRps4 recognition occurs through two RRS1 protomers found in a heterotetramer with two RPS4 protomers^43^. Following recognition of AvrRps4, the RPS4 TIR domains are activated to produce signaling molecules. Production of 2ʹcADPR leads to activation of EDS1-PAD4-mediated immunity, while production of ADPR-ATP activates EDS1-SAG101^4,7^. **C-E.** Proliferation of *P. syringae* pv. tomato DC3000 Δ*hopQ1* strains carrying operons expressing the indicated proteins in leaves of WT *A. thaliana* Col-0 (**C**), *pad4-1* mutant (**D**), or *sag101-3* mutant (**E**). HopQ^DD^ denotes the catalytically inactive HopQ^D105N, D384N^. Shown are colony-forming units (CFUs) per cm^2^, measured four days post inoculation from three independent experiments with four replicates each (*n* = 12 in total). Data are presented as box plots: median (center line), interquartile range (IQR, box), and the most extreme values within 1.5 x IQR (whiskers). Individual measurements are shown as dots. Pairwise, two-sided Mann-Whitney U tests followed by Benjamini-Hochberg correction were used to assess statistical significance of differences in bacterial proliferation between strains within each plant genotype. ns, not significant.

It was recently shown that the RPS4 TIR domain produces two structurally distinct classes of signaling molecules. In addition to cyclizing ADPR into 2ʹcADPR^7^, the RPS4 TIR can attach ADPR to another molecule, generating ADPR conjugated to ATP (ADPR-ATP) or to a second ADPR (di-ADPR). These linear ADPR conjugates were found to bind and activate the EDS1-SAG101 immune protein complex^4^. Thus, it is currently hypothesized that the same plant TIR-domain protein can simultaneously activate two distinct EDS1 immune nodes^1,45^: one that relies on 2ʹcADPR which, once converted to pRib-AMP, activates EDS1-PAD4^7^, and another in which ADPR conjugates directly activate EDS1-SAG101^4^ (Figure 5B). Consistent with this model, it was previously shown that deficiency of either PAD4 or SAG101 node renders *A. thaliana* more susceptible to infection by a *P. syringae* encoding *avrRps4*^38^. As HopQ1 is specific for 2ʹcADPR, we reasoned that the failure of HopQ1-expressing bacteria to overcome AvrRps4-triggered immunity in WT plants reflects the activity of the EDS1-SAG101 immune node that functions in parallel to the EDS1-PAD4 one^46,47^.

To examine this hypothesis, we spray-inoculated the *P. syringae* strains tested above onto leaves of *A. thaliana* mutant plants lacking either PAD4 or SAG101. Strikingly, when bacterial strains expressing AvrRps4 were spray-inoculated onto plants lacking SAG101, the WT HopQ1 conferred ∼300-fold increase in bacterial proliferation compared to the catalytically inactive HopQ1 variant (Figure 5D; Figure S9B; Table S5). These results demonstrate that the catalytic activity of HopQ1 specifically suppresses AvrRps4-triggered EDS1-PAD4 immunity. In plants lacking PAD4, however, proliferation of AvrRps4-expressing bacteria remained restricted regardless of whether the strains expressed WT or catalytically inactive HopQ1 (Figure 5E; Figure S9C; Table S5). Together, these data reveal that HopQ1 can efficiently inhibit TIR-induced, PAD4-dependent immunity to promote bacterial virulence.

## Discussion

XopQ was discovered more than two decades ago as a protein delivered into plant cells by several plant pathogenic bacteria^20,23,24^, but although XopQ homologs have since been identified across diverse lineages of plant pathogens, the molecular mechanism of its contribution to virulence has remained elusive. In this work we show that XopQ is an enzyme that degrades the TIR-derived signaling molecule 2ʹcADPR, thus inhibiting 2ʹcADPR-dependent plant immunity. The function of XopQ is analogous to the function of multiple families of phage proteins that were shown to inhibit bacterial immunity by cleaving or sequestering TIR-derived immune signals^13–16,19^. Human immune signaling molecules such as cyclic GMP-AMP (cGAMP) and cyclic oligoadenylates (cOA) were also shown to be targeted for cleavage by dedicated viral proteins as a means of countering immunity^48^. Our results now show that cleavage of immune signaling molecules is a common host takeover strategy conserved across pathogens of bacteria, humans and plants.

XopQ was originally classified as a nucleoside hydrolase based on predicted structural homology^20^. Subsequent structural studies supported this annotation but also revealed that the XopQ active site is larger than that of canonical nucleoside hydrolases, which led to the suggestion that this enzyme acts on a “novel” substrate^28,29^. Indeed, 2ʹcADPR – a molecule that had not yet been discovered when the XopQ structures were reported – is approximately twofold larger than a canonical nucleoside (541 Da versus ∼270 Da, respectively). XopQ is further distinguished from known nucleoside hydrolases by the chemical bond it hydrolyzes. Whereas canonical nucleoside hydrolases cleave the *N*-glycosidic bond linking a nitrogenous base to a ribose, XopQ hydrolyzes the *O*-glycosidic bond connecting the two ribose moieties of 2ʹcADPR while leaving the adenine-ribose *N*-glycosidic bond intact. Moreover, XopQ displays a catalytic efficiency higher than that reported for almost all characterized nucleoside hydrolases. Together, these observations point to a substantial evolutionary divergence of XopQ from canonical members of this enzyme family. In this regard, XopQ joins a growing list of immune-related enzymes, including NAD reconstitution enzymes^49^ and purine nucleoside phosphorylase (PNP)^50^, whose catalytic activities appear to have been remodeled in the process of adaptation from housekeeping roles to functions in host-pathogen interactions. These examples illustrate how evolutionary arms races can drive the emergence of novel catalytic functions.

The widespread use of XopQ by bacteria that infect a wide array of flowering plants, together with the conserved catalytic activity we demonstrate in this study, points to 2ʹcADPR as a central signaling molecule in plant immunity. Consistent with this notion, 2ʹcADPR appears as the predominant product of many plant TIR proteins examined to date^2,3,51,52^. Nevertheless, how 2ʹcADPR promotes immunity has remained ambiguous. In a recent study, infiltrating *A. thaliana* leaves with 2ʹcADPR stimulated PAD4-dependent immune responses, however pRib-AMP instead of 2ʹcADPR was detected bound to the EDS1-PAD4 complex^7^. It was therefore suggested that 2ʹcADPR is converted in plant cells to pRib-AMP, which then directly activates the EDS1-PAD4 immune node. Our data support this hypothesis, showing that pRib-AMP appears in *N. benthamiana* leaves following BdTIR-mediated 2ʹcADPR production, and that XopQ-mediated depletion of 2ʹcADPR results in a concomitant elimination of pRib-AMP (Figure 4). Together with our findings that XopQ does not cleave pRib-AMP as a substrate, and that HopQ1, the *P. syringae* XopQ homolog, suppresses PAD4-dependent immunity, our observations corroborate a model in which TIR-produced 2ʹcADPR is converted into pRib-AMP to activate the EDS1-PAD4 immune complex.

The current view of the plant-pathogen arms race, originally formulated as the zigzag model^53^, proposes a sequence of immunity and counter-immunity evolutionary events. In this model, intracellular immune receptors, including TIR-containing proteins, evolved as a second layer of defense to recognize effectors secreted by pathogens to suppress basal immunity initiated by cell-surface receptors. XopQ represents a pathogen counter-adaptation to this second immune layer, likely emerging in response to the extensive deployment of TIR proteins in plant immunity. Remarkably, the *Nicotiana* lineage evolved the intracellular receptor Roq1, which directly recognizes XopQ as a signature for infection^54^. Upon XopQ recognition, the TIR domain of Roq1 becomes catalytically active^55^, and intriguingly produces 2ʹcADPR^3,51^, raising an apparent mechanistic paradox: the immune receptor that detects XopQ uses the very same signaling molecule that XopQ can inactivate. A possible resolution to this paradox comes from a structural study that examined Roq1 in complex with XopQ^55^, showing that Roq1 interacts with XopQ in a way that likely physically blocks access of 2ʹcADPR to the XopQ active site, and therefore presumably inhibits enzymatic activity^55^. As XopQ targets a fundamental node of plant immunity and is widely distributed among plant pathogenic bacteria, it is likely that plants have evolved additional mechanisms to counteract its activity.

Extending this evolutionary perspective, it is possible that XopQ and other pathogen-encoded inhibitors of 2ʹcADPR signaling have led to the evolution of the EDS1-SAG101 immune node in dicotyledonous plants^56^, which is activated by molecules other than 2ʹcADPR or its derivatives. Following this logic, additional 2ʹcADPR-independent TIR signaling pathways may exist in monocotyledonous plants, as this lineage employs PAD4 but naturally lacks SAG101, and is routinely infected by XopQ-encoding pathogens. As direct targeting of immune signaling molecules appears to be a recurring strategy utilized by pathogens across the tree of life, virulence mechanisms that target ADPR-ATP, di-ADPR, or other plant TIR-derived immune activators likely also exist and remain to be discovered.

## Methods

### Bacterial strains

*E. coli* MG1655 and LOBSTR-BL21 (DE3)^57^ strains were grown in magnesium manganese broth (MMB; LB + 0.1 mM MnCl_2_ + 5 mM MgCl_2_) at 37 °C or 16 °C with shaking at 200 RPM. *A. tumefaciens* GV3101 (pMP90) strains were grown in LB at 28 °C with shaking at 250 RPM. *P. syringae* pv. tomato DC3000 strains where grown in NYGA 28 °C. Whenever required, the appropriate antibiotics were added in the following concentrations: spectinomycin (50 μg ml^−1^ or 100 μg ml^−1^ for *E. coli* or *A. tumefaciens*, respectively, ampicillin (100 μg ml^−1^), kanamycin (50 μg ml^−1^), gentamycin (25 μg ml^−1^ or 10 μg ml^−1^ for *A. tumefaciens* or *P. syringae*, respectively), rifampicin (50 μg ml^−1^).

BdTIR under the control of a tetracycline inducible promoter was integrated into the *E. coli* MG1655 genome downstream of the glmS gene using Tn7 integration plasmid as previously described^16^. For expression in *E. coli* K-12 MG1655, XopQ proteins were cloned and transformed as previously described^58^. Briefly, XopQ or GFP genes were cloned under the control of an IPTG inducible promoter into the thrC-Phspank vector, then introduced to *E. coli* K-12 MG1655 strains, in which either BdTIR or the Type VI Thoeris defense system (from metagenomic, Gammaproteobacteria scaffold)^12^ were integrated into the bacterial genome.

To generate *E. coli* LOBSTR-BL21 (DE3) strains, XopQ homologs were cloned downstream of the N-terminal His14-bdSUMO tag, and under the control of an IPTG inducible promoter using the pET28-His-bdSUMO expression vector^59^.

To generate *A. tumefaciens* GV3101 strains, genes were cloned downstream of the 35s promoter using the pH2GW7 expression vector^60^, replacing the entire gateway cassette.

To generate *P. syringae* pv. tomato DC3000 strains, operons were cloned into the pCK013 vector^61^, replacing the entire gateway cassette and the downstream Myc-tag encoding sequences. The operons were under the regulation of *hopQ1* native promoter and terminator. The two open reading frames in the operons were separated by the native ribosome binding site, defined as 20 nucleotides upstream to the first codon of *avrRps4* as in *P. syringae* pv. pisi.

All plasmids were synthesized by Twist Bioscience or GenScript Corporation, after codon optimization to the appropriate host. Transformations into *E. coli*, *A. tumefaciens*, and *P. syringae* were performed using standard electroporation. All gene accessions and protein sequences used in this study are listed in Table S1. Operon sequences used in the experiments shown in Figure 5 are in Table S4.

### Phage strains

The *E. coli* phage SECphi18 (GenBank: LT960609.1), previously isolated by us^9^, was propagated on *E. coli* MG1655 or by picking a single phage plaque into a liquid culture grown at 37 °C to an optical density at 600 nm (OD_600_) of 0.3 in MMB broth until culture collapses. The culture was then centrifuged for 10 minutes at 3,200 x *g* and the supernatant was filtered through a 0.2 µm filter to get rid of remaining bacteria and bacterial debris.

### AlphaFold structural predictions

AlphaFold3 with default parameters was used to generate structural models for XopQ proteins together with the respective ligands. The ligands SMILES codes are listed in Table S6. Structural models were visualized using the PyMOL Molecular Graphics System, version 3.1.3.1.

### XopQ functional characterization in *E. coli*

*E. coli* MG1655 strains carrying BdTIR integrated in the genome downstream of the glmS gene and expressing the indicated XopQ constructs from the thrC-Phspank vector were grown in 50 ml of liquid MMB. When the cultures reached an OD_600_ of 0.3, IPTG was added to a final concentration of 1mM to induce XopQ expression. Cultures were then incubated until reaching an OD_600_ of 0.6, at which point BdTIR expression was induced by addition of anhydrotetracycline to a final concentration of 100 nM. Cultures were further grown for 1.5 hours or until reaching an OD_600_ of 1.

Cell harvested by centrifugation at 3,200 x *g* for 10 minutes at 4 °C and resuspended in a lysis buffer (20 mM HEPES pH 7.3, 125 mM MgCl_2_) at a ratio of 600 µl per culture normalized to OD_600_ = 1. Cells were lysed using Lysing Matrix B in a bead beater (two cycles of 6 m/s for 40 seconds). Lysates were clarified by centrifugation at 16,000 x *g* for 10 minutes at 4 °C and filtered with 3 kDa Amicon filters by centrifugation at 12,000 x *g* for 45 minutes at 4 °C. Filtered lysates were stored at -20 °C until HLPC-MS analysis.

### LC-PDA and LC-PDA-MS of bacterial lysates and enzymatic reactions

The samples shown in Figure 1D were analyzed on Agilent 1260 HPLC system coupled with Thermo Fisher Diode Array Detector. The analytes were separated by a gradient elution with SUPELCOSIL™ LC-18-T HPLC Column. Mobile phase A consisted of 20 mM potassium phosphate pH 6 and mobile phase B consisted of 20 mM potassium phosphate pH 6 in 20% methanol. The gradient was as follows: 2 minutes of mobile phase A 100%, 5 minutes 75% A and 25% B, 2 minutes 50% A and 50% B, gradual increase over 2 minutes to 100% A, 3 minutes 100% A, and 5 minutes 100% B, at 0.8 ml/minutes flow rate. The Diode Array detector was set to measure UV absorbance at 260 nm.

All others bacterial lysate samples, were carried out on Waters Acquity UPLC system (Manchester, UK) coupled with Waters Photo-Diode-Array (PDA) and QDa detectors. The analytes were separated by gradient elution with Waters Acquity Premier HSS T3 Column, 1.8 μm, 2.1 × 100 mm at 0.4 ml/minutes flow rate, 35°C. Mobile phase A consisted of 20 mM aqueous Ammonium Acetate (Sigma-Aldrich 09691) and mobile phase B consisted of 20 mM Ammonium Acetate in acetonitrile:water ratio of 75:25. The gradient was as follows: 0–2.5 minutes 0% of B, 2.5-6 minutes linear gradient ramp to 10% of B, 6-6.1 minutes ramp to 100% of B. PDA was set to measure UV absorbance between 230 and 330 nm. QDa with electro spray ionization source was operated in negative mode (ESI-) and source temperature was set at 140°C. Single ion monitoring parameters were set as following: 2ʹ3ʹcAMP: selected ion recording (SIR) m/z 328.00, cone voltage 20 V. 2ʹ3ʹcGMP: SIR m/z 344.08, cone voltage 20 V. cADPR isomers: SIR m/z 540.05, cone voltage 25 V. ADPR and pRib-AMP: SIR m/z 558.06, cone voltage 20 V.

### Protein purification

*E. coli* LOBSTR-BL21 (DE3) strains expressing the indicated XopQ construct from the pET28-His-bdSUMO vector were grown in 500 ml MMB at 37 °C until the culture reached an OD_600_ of 0.6. IPTG was then added to a final concentration of 1mM to induce expression of His-bdSUMO-XopQ, and cultures were incubated for ∼12 hours at 16 °C.

Cells were harvested by centrifugation at 4,000 x *g* for 15 minutes at 4 °C, resuspended in 50 ml lysis buffer (20 mM HEPES pH 7.3, 0.4 M NaCl, 1 mM DTT, 30 mM imidazole), lysed using the EmulsiFlex-C3 homogenizer (Avestin, Canada) and clarified by centrifugation at 20,000 x *g* for 20 minutes at 4 °C, followed by filtration. Clarified lysates were applied to a 5 ml His-Trap FF column (Cytiva), pre-equilibrated with lysis buffer. The column was washed with lysis buffer, followed by a high-salt wash (1 M NaCl), and proteins were eluted with buffer containing 20 mM HEPES (pH 7.3), 0.4 M NaCl, 1 mM DTT, and 300 mM imidazole.

Eluted proteins (∼15 ml) were dialyzed overnight at 4 °C in 3.5 kDa MWCO Slide-A-Lyzer™ cassettes (Thermo Scientific), in cleavage buffer (20 mM HEPES, 125 mM KCl, 1 mM DTT) with 1 μM His-tagged bdSENP1 protease. Following cleavage, samples were supplemented with NaCl and imidazole to final concentrations of 275 mM and 40 mM, respectively, and reloaded onto a HisTrap FF column to remove uncleaved protein, the cleaved bdSUMO tag, and the bdSENP1 protease. The flow-through, containing the cleaved, untagged XopQ protein, was collected, aliquoted and stored at -80 °C. Protein concentration was determined using the Qubit^TM^ protein assay kit.

### Enzymatic reactions

XopQ catalytic activity was assessed by incubating the indicated protein-substrate combinations at final concentrations of ∼100 nM (protein) and 10 µM (substrate) in a reaction buffer (20 mM HEPES pH 7.3, 125 mM MgCl_2_) for 20 minutes at 25 °C. Reactions were filtered using 3 kDa Amicon filters by centrifugation at 12,000 x *g* for 45 minutes at 4 °C, and stored at -20 °C until HPLC-MS analysis.

To assess the effect of increasing 3ʹcADPR concentrations on XopQ-catalyzed 2ʹcADPR hydrolysis, reactions were performed with ∼10 nM XopQ, 1 µM 2ʹcADPR and 3ʹcADPR at 10 µM or 1 mM. Reactions were incubated for 10 minutes at 25 °C in reaction buffer (20 mM HEPES pH 7.3, 125 mM MgCl_2_) and terminated by heating at 95 °C for 5 minutes. Samples were then centrifuged and stored at -20 °C until HPLC-MS analysis.

To assess the kinetics of XopQ-catalyzed 2ʹcADPR hydrolysis, reactions were performed with ∼0.05 nM XopQ and 2ʹcADPR at increasing concentrations (75-1250 nM) and incubated for 2 minutes at 25 °C. Reactions were terminated by heating at 95 °C for 5 minutes, after which samples were centrifuged and stored at -20 °C until HPLC analysis. ADPR production rates were calculated using a pre-defined ADPR standard curve based on UV absorbance in UPLC–PDA-QDa. Kinetic parameters (*k_cat_* = 4.805 s^-1^, 95% CI 4.1 - 5.4 s^-1^; *K*_m_ = 336 nM, 95% CI 231–441 nM; *k_cat_*/*K*_m_ = 1.429 X 10^7^ M^-1^ s^-1^, 95% CI 1.046 X 10^7^ - 1.813 X 10^7^ M^-1^ s^-1^) were obtained by weighted nonlinear least-squares regression to a Michaelis-Menten model, using the curve_fit function from the scipy.optimize module in Python, with standard error of the mean used as weights.

To compare the catalytic properties of XopQ with those of other enzymes belonging to the “inosine-uridine preferring nucleoside hydrolases” (Pfam accession: PF01156), kinetic parameters for enzymes assigned to the following Enzyme Commission (EC) classes were retrieved from the BRENDA database^35^: EC.3.2.2.1, EC.3.2.2.2, EC.3.2.2.3, EC.3.2.2.7, EC.3.2.2.8, EC.3.2.2.13, EC.3.2.2.25.

### Plaque assay

Phage titers were determined using the small-drop plaque assay^62^. An overnight culture of *E. coli* (300 µL) was mixed with 30 ml MMB containing 0.5% agar and the appropriate inducers (1 mM IPTG for XopQ expression and 100 nM anhydrotetracycline for type VI Thoeris), poured onto 10 cm square plates, and allowed to solidify for 1 hour at room temperature.

Tenfold serial dilutions of SECphi18 coliphage were prepared in MMB, and 10 µL drops were applied onto the agar surface. After the drops had dried, plates were inverted and incubated overnight at 25 °C. Plaque-forming units (PFUs) were determined by counting the plaques after overnight incubation.

The *E. coli* strains used in this assay included a negative control strain containing genomically integrated RFP and a thrC-Phspank plasmid carrying GFP, as well as strains containing genomically integrated type VI Thoeris, either without a plasmid or expressing XopQ or XopQ^D120N^ from the thrC-Phspank plasmid.

### Infection in liquid culture

Overnight cultures of bacteria were diluted 1:100 in MMB medium containing the appropriate inducers (1 mM IPTG for XopQ expression and 100 nM anhydrotetracycline for type VI Thoeris) and incubated at 37 °C with shaking at 200 RPM. When cultures reached optical density at OD_600_ of 0.3, cells were transferred to a 96-well plate (180 µL per well) and infected with 20 µL of the SECphi18 coliphage at the indicated multiplicities of infection. Plates were incubated at 25 °C with shaking in a TECAN Infinite 200 plate reader, and OD_600_ was monitored every 10 minutes.

### 2ʹcADPR-induced toxicity assay

Overnight cultures of bacteria were diluted 1:50 in MC medium (80 mM K_2_HPO_4_, 30 mM KH_2_PO_4_, 2% glucose, 30 mM trisodium citrate, 22 μg/ml ferric ammonium citrate, 0.1% casein hydrolysate, 0.2% potassium glutamate) containing the appropriate inducers (1 mM IPTG for XopQ expression and 100 nM anhydrotetracycline for type VI Thoeris) and incubated at 37 °C with shaking at 200 RPM.

When cultures reached OD_600_ of 0.3, cells were transferred to a 384-well plate (18 µL per well) and mixed with 2ʹcADPR (2 µL per well) to achieve a final concentration of 1 mM. Plates were incubated at 37 °C with shaking in a TECAN Infinite 200 plate reader, and OD_600_ was monitored every 10 minutes.

### Analysis of Thoeris systems in *Pseudomonadota*

Bacterial genomes (Table S3) were retrieved from the NCBI database. DefenseFinder^63^, supplemented with models to include Thoeris types V-XI as previously described^12^, was applied to determine the defense-system repertoire of each genome. Genomes were screened for XopQ homologs using BLAST (minimum coverage of 80%, minimum identity of 45%, and an expected E-value < 1×10^-5^), with the six XopQ homologs listed in Table S3 as queries.

### Plant growth

*N. benthamiana eds1 pad4 sag101a sag101b* (*Nb epss*) plants were grown as previously described^64^. Seeds were sown in 0.25 L plastic pots filled with a peat-based greenhouse substrate and covered with granulated cork to prevent pest infestation. Plants were cultivated in a greenhouse cabin exposed to ambient sunlight and supplemented with a broad-spectrum 16 h photoperiod from LED lights. Pots were daily irrigated with a nutrient solution containing an electrical conductivity of 2.2 mS/cm; pH 5.6; potassium (0.46 mmol/L); calcium (0.38 mmol/L); magnesium (0.16 mmol/L); nitrogen-to-potassium ratio of 1.8; ammonium fraction of total nitrogen of 0.05 and phosphorus ratio of 0.055.

*A. thaliana* Col-0, *pad4-1*, and *sag101-3* seeds were sown in 0.25 L plastic pots filled with a peat-based greenhouse substrate and covered with granulated cork to prevent pest infestation. Plants were cultivated in a growing chamber under conditions of 10 hours light and 14 hours dark cycles, with a light intensity of approximately 150 μmol/m²/s, at 22 °C and 65 % humidity.

### XopQ functional characterization in *N. benthamiana*

Overnight cultures (10 ml) of *A. tumefaciens* strains grown in LB with the appropriate antibiotics were centrifuged for 6 minutes at 3,200 x *g*, resuspended in 5 ml infiltration buffer (10 mM MgCl_2_, 10 mM MES pH 5.6, and 250 µM acetosyringone), and diluted to OD_600_ of 1.

Cells at OD_600_ of 1 were infiltrated into *N. benthamiana* to assess expression of individual proteins by western blot analysis.

For co-infiltration of two *A. tumefaciens* strains, (e.g., BdTIR and XopQ), cultures were mixed at a 1:1 ratio, resulting in an OD_600_ of 0.5 for each strain and a total OD_600_ of 1.

Co-infiltrations were performed across the entire area of the first three leaves of ∼3-week-old plants, using four plants per gene combination. Plants were kept in the dark for ∼12 hours following infiltration and then returned to the greenhouse. At 48 hours post infiltration, infiltrated leaves were collected into four samples, each comprising pooled material from different plants, flash-frozen in liquid nitrogen, and ground using a pre-chilled ceramic mortar and pestle under frozen conditions.

Metabolites were extracted from 200 mg of tissue per sample using pre-cooled 50% Methanol:H2O for 30 minutes at 4 °C with shaking at 1,500 RPM. Samples were then centrifuged at 21,000 x *g* for 10 minutes at 4 °C, and the clarified supernatants were evaporated to dryness using a Speed Vac concentrator (Centrivap; Labconco) at 10°C at 1,000 RPM and stored at -80 °C until Ion-chromatography (IC)-MS analysis.

### IC-MS analysis samples derived from plant extracts

Metabolite separation was performed with a Dionex Integrion HPIC System (Thermo Fisher) equipped with an IonPac AS11-HC column (2 mm × 250 mm, Thermo Fisher) maintained at 30 °C. A guard column, (IonPac AG11-HC b, 2 mm × 50 mm, Thermo Fisher), was installed upstream of the separation column. The eluent (KOH) was generated in situ by a KOH cartridge and deionized water.

Separations were carried out at a flow rate of 0.380 ml/minutes using the following gradient: 0–2 minutes 35 mM KOH, 2-6 minutes 35-100 mM, 6-8 minutes 100 mM, 8,01-13 minutes 35 mM. A Dionex suppressor AERS 500 (2 mm) was used for the exchange of the KOH. The suppressor pump flow was set at 0.6 ml/minutes.

Samples were re-dissolved in 90 µL LC-MS-grade water and injected at 8 °C using full-loop mode (5 µL). The Integrion system was coupled to a Q Exactive mass spectrometer operating in negative ionization mode with full MS scanning over an m/z range of 59-602. Parallel reaction monitoring (PRM) was performed for 2ʹcADPR (ms2 540.0538 at Higher-energy Collisional Dissociation (HCD) of 35.00 [50.0000-570.0000]) and ADPR/pRib-AMP (ms2 558.0644 at HCD of 35.00 [50.0000-590.0000]). The resolution was set to 17,500, and the maximal injection time was set to 200 msec. The HESI source was operated with a spray voltage of 2.75 kV. The ion transfer capillary temperature was 350 °C, the sheath gas flow was 50 arbitrary units (AU), the auxiliary gas flow 14 AU, and the sweep gas flow was set to 3 AU at 380 °C.

### Western blot analysis

Western blot analysis of protein expression in *N. benthamiana* was performed as previously described^64^. In summary, 100 mg of transformed leaf tissue per sample was flash-frozen and pulverized using a bead beater. The frozen powder was resuspended in lysis buffer (50 mM Tris-HCl, pH 7.4, 150 mM NaCl, 5% glycerol, 10 mM DTT, 0.5% polysorbate 20, 1x Protease inhibitor mix P (Serva), 5% BioLock (IBA Lifesciences); pH adjusted to 7.4) and centrifuged twice at 16,000 x *g*. The clarified supernatant was supplemented with 4x Lämmli buffer containing 5% mM β-mercaptoethanol and heated for 5 minutes at 95 °C, followed by cooling on ice.

Samples (10 µL each) were resolved on a 10% SDS-PAGE gel and transferred to a PVDF membrane. Membranes were blocked in TBS-T containing 5% (w/v) milk for 1 hour at room temperature (RT), washed three times for 5 minutes in TBS-T, and incubated with anti-HA antibody (1:2,000) in TBS-T containing 5% BSA for 1 hour at RT. Membranes were then washed three times for 10 minutes in TBS-T and incubated with secondary anti-rabbit antibody (1:2,000) in TBS-T containing 5% milk for 1 hour at RT. After three additional washes (15 minutes each) in TBS-T, signals were developed using SuperSignal West Femto substrate.

### Pseudomonas syringae infection

Five-week-old *A. thaliana* plants were covered before infection for 3–4 h with a transparent lid pre-sprayed with water. Plants were subsequently spray-inoculated with the *Pseudomonas syringae* pv. tomato Δ*hopQ1* strains adjusted to an OD_600_ of 0.2 in a 10 mM MgCl₂ solution containing 0.04% (v/v) Silwet L-77. Silwet L-77 was added immediately prior to inoculation to minimize potential detrimental effects on bacterial viability. Spray inoculation was performed using a hand sprayer until all leaves were uniformly wetted and runoff was observed, ensuring coverage of both adaxial and abaxial leaf surfaces. Following inoculation and dry out of the bacteria solution, plants were transferred to an infection growth chamber maintained under an 8 hours light/16 hours dark photoperiod at 23 °C and 60% relative humidity, and were covered with a transparent lid for 48 hours to maintain high humidity conditions. For bacterial growth quantification, samples designated as day 0 were harvested 1 hour after spray inoculation, whereas bacterial growth was assessed at 4 days post inoculation (dpi). Bacteria were recovered from leaf discs collected from infected plants (three leaf discs per sample, each obtained from a different leaf; 0.6 cm diameter). Leaf discs were incubated in either 1 ml (0 dpi samples) or 1.5 ml (4 dpi samples) of 10 mM MgCl₂ solution containing 0.01% Silwet L-77 and shaken at 650 RPM, 28 °C for 1 h. Bacterial suspensions were then serially diluted and plated onto selective NYGA medium. Colony-forming units (CFUs) were quantified after incubation at 28 °C for 2 days.

## Acknowledgements

We thank members of the Sorek lab, the Schulze-Lefert lab, and the Parker lab for constructive discussions during this study. We also thank Dr. Alla H. Falkovich for technical assistance, Dr. Doron Teper for kindly providing the WT and Δ*hopQ1 Pseudomonas syringae* pv. tomato DC3000 strains, and Prof. Asaf Aharony for assistance with plant-growing facilities. R.S. was supported, in part, by the European Research Council (grant ERC-AdG GA 101018520), the Israel Science Foundation (MAPATS grant 2720/22), the Deutsche Forschungsgemeinschaft (SPP 2330, grant 464312965), the Minerva Foundation with funding from the Federal German Ministry for Education and Research, and a research grant from Magnus Konow in honor of his mother Olga Konow Rappaport. O.R. was supported by a Weizmann Postdoctoral Excellence Fellowship. P.S.L, J.E.P., A.W.L, S.P., E.L. and F.L. were supported by the Max-Planck Society. F.L. and J.E.P. were also supported by grants within DFG CRC-1403 project 414786233 and iHEAD (NRW Profilbildung ID: PB22-025A). Figures 4A and 5A were generated with icons exported from BioRender.com. R.S. is a scientific cofounder and advisor of Ecophage.

## Supplementary Figures

**Figure S1.**
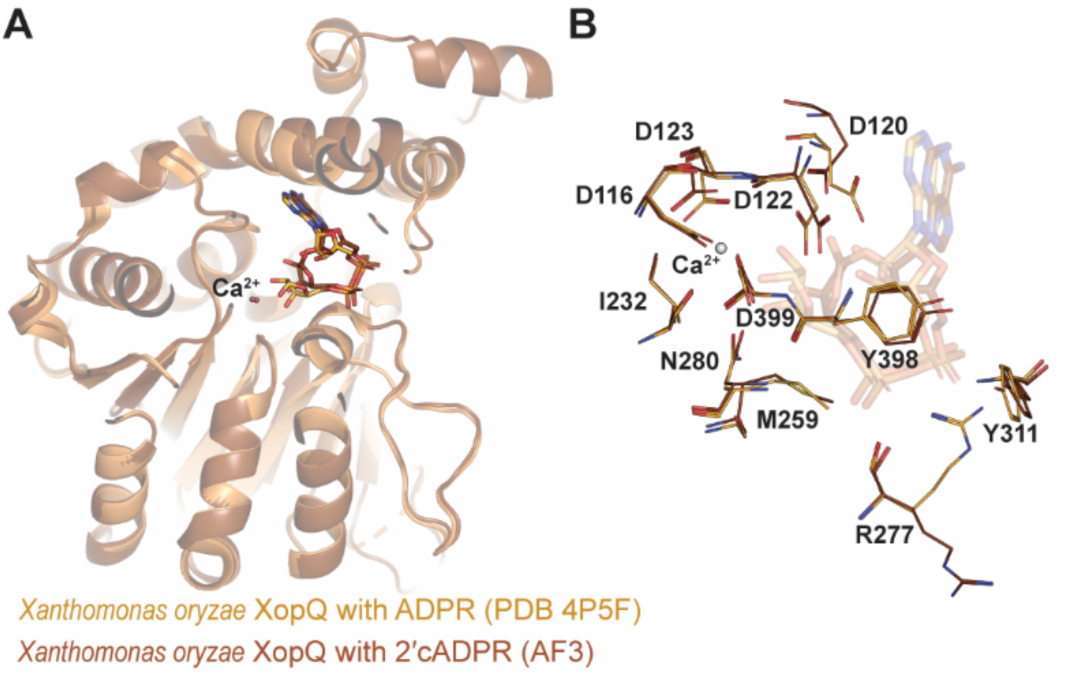
Structural comparison of XopQ bound to ADPR and the predicted binding to 2ʹcADPR. **A.** Superimposition of the crystal structure of XopQ bound to ADPR (PDB 4P5F) and the AlphaFold3 model of *X. oryzae* XopQ bound to 2ʹcADPR and Ca^2+^. **B.** XopQ residues located within 3 Å of ADPR in the crystal structure, shown superimposed with the same residues in the AlphaFold3 model with the 2ʹcADPR ligand. Residues from the crystal structure (including the ADPR ligand) are in yellow, those from the AlphaFold3 model are in brown.

**Figure S2.**
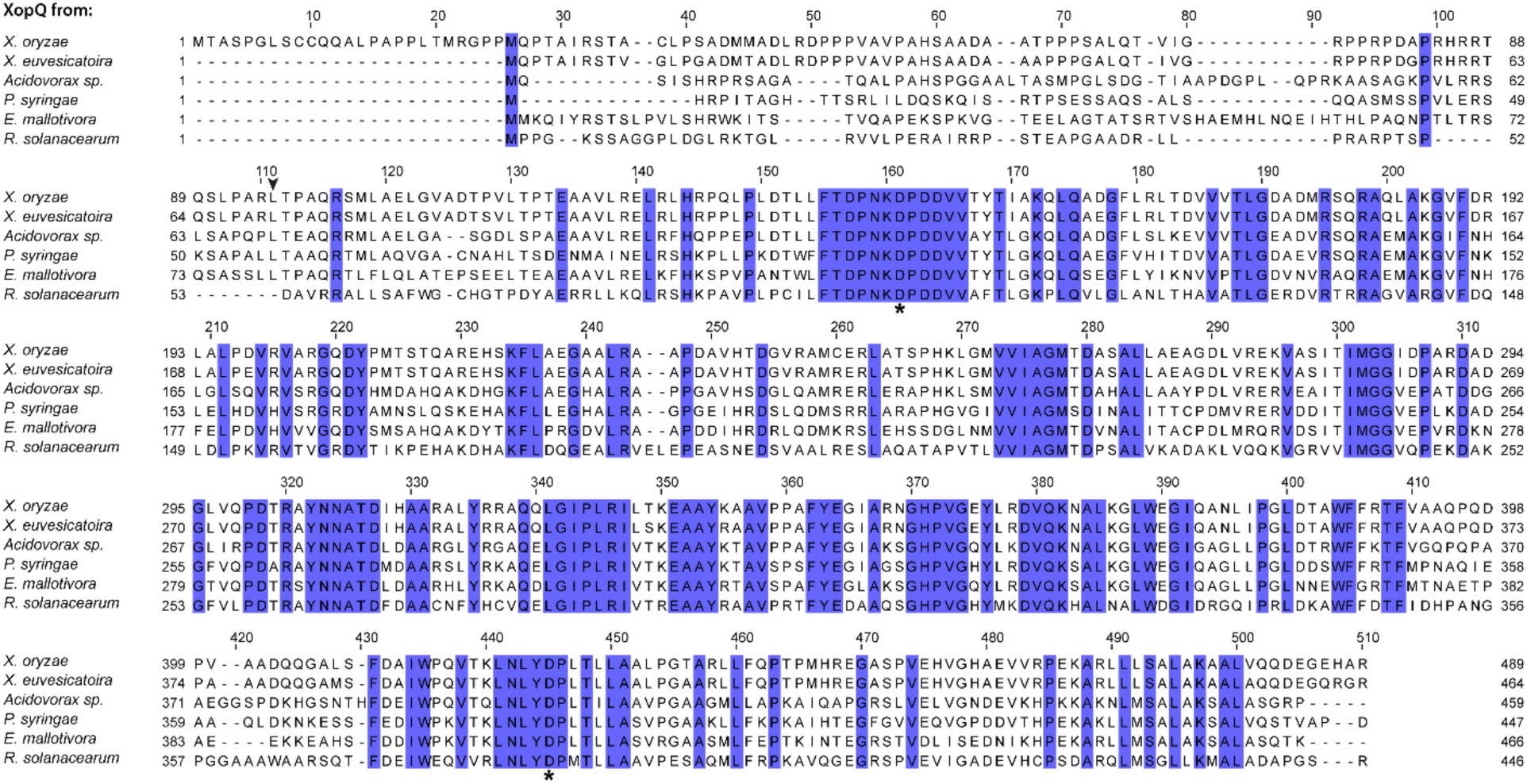
Multiple sequence alignment of the six XopQ proteins used in this study. Conserved residues (100%) are highlighted in blue. Arrowhead marks the end of the unstructured N-terminus that was truncated from the XopQ proteins used for the in vitro assays. Asterisks indicate the conserved aspartic acid residues mutated in this study. The alignment was generated using MAFFT^65^ with default parameters.

**Figure S3.**
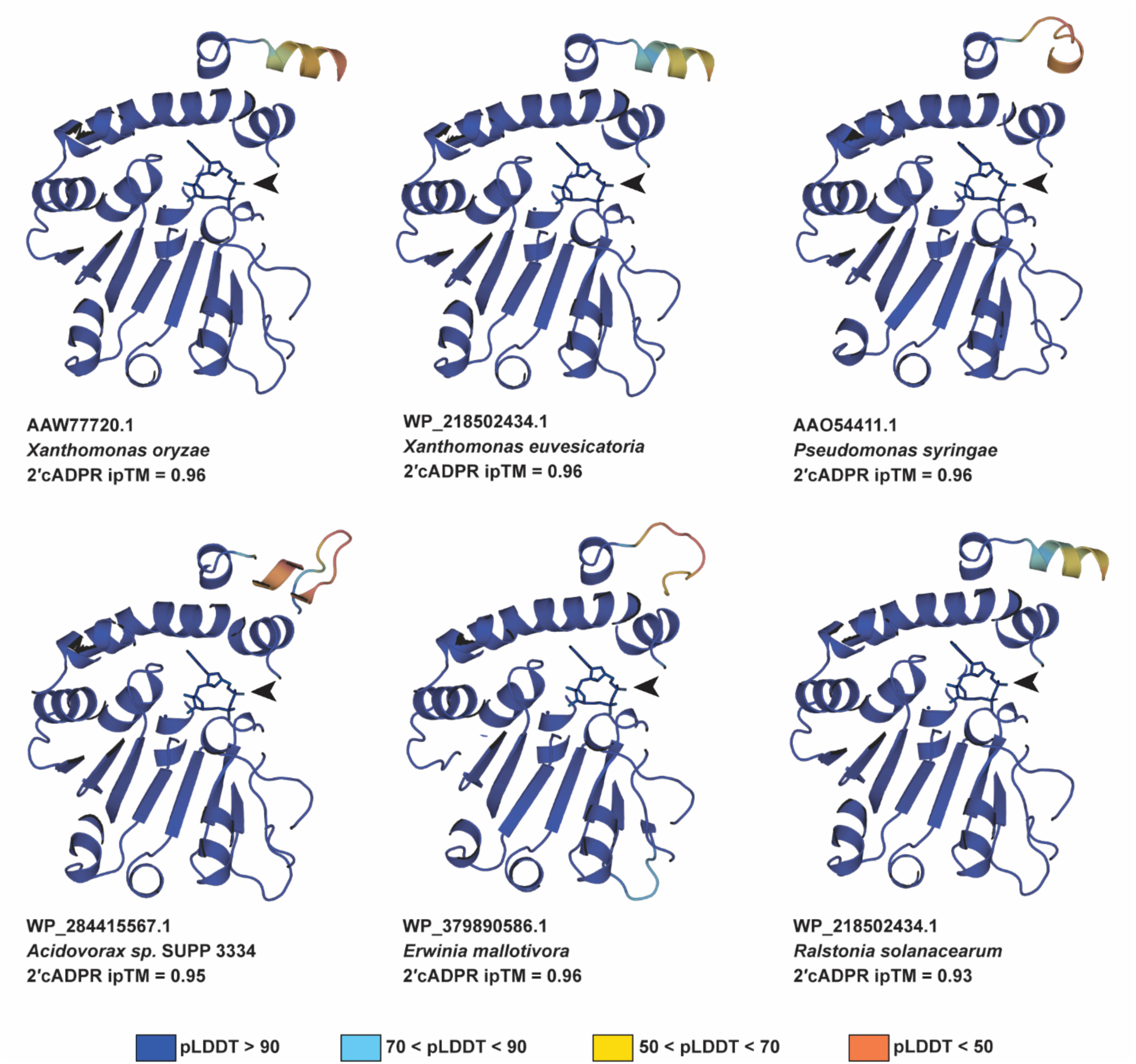
AlphaFold3 models for XopQ homologs bound to 2ʹcADPR. Shown are AlphaFold3 models for XopQ homologs co-folded with 2ʹcADPR and a Ca^2+^ ion. Arrowheads indicate the 2ʹcADPR molecules. NCBI accession number is indicated for each protein. Colors represent pLDDT scores. Presented ipTM values are the average of 5 AlphaFold3 models.

**Figure S4.**
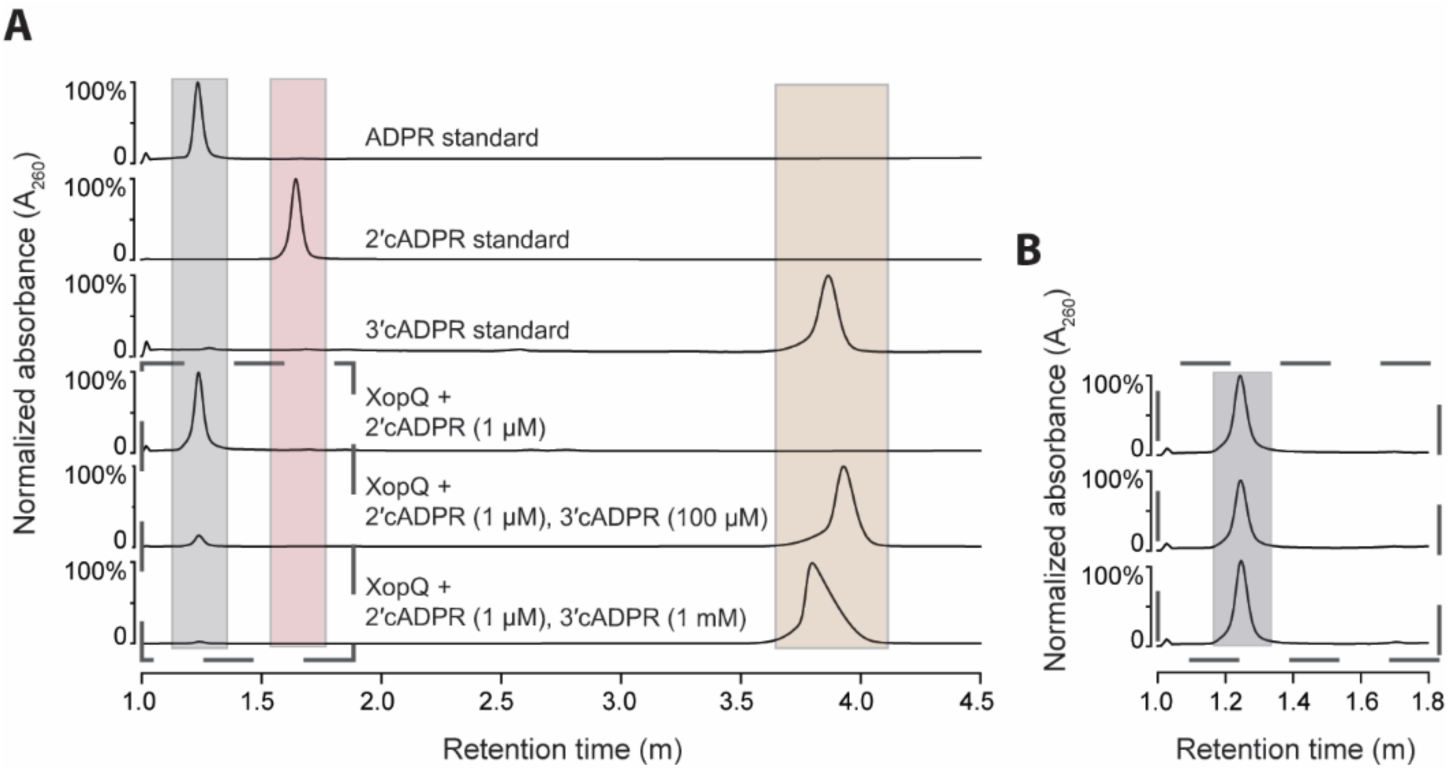
Complete hydrolysis of 2ʹcADPR to ADPR by XopQ at increasing 3ʹcADPR-to-2ʹcADPR molar ratios. **A.** HPLC chromatograms of in vitro reactions examining the catalytic activity of *X. euvesicatoria* XopQ toward 2ʹcADPR (1 µM) in the presence of 100 µM or 1 mM 3ʹcADPR. Data are presented as absorbance values at A_260_, with each chromatogram normalized independently to its maximum value. Shading represents retention time of ADPR (grey), 2ʹcADPR (pink), and 3ʹcADPR (brown). **B.** HPLC chromatograms showing similar ADPR levels for the three reactions shown in the dashed box in panel A. Data are presented as absorbance at A_260_. Absorbance values were normalized to the reaction containing XopQ and 1 µM 2ʹcADPR. Grey shading represents retention time of ADPR. Shown are representatives of at least three replicates.

**Figure S5.**
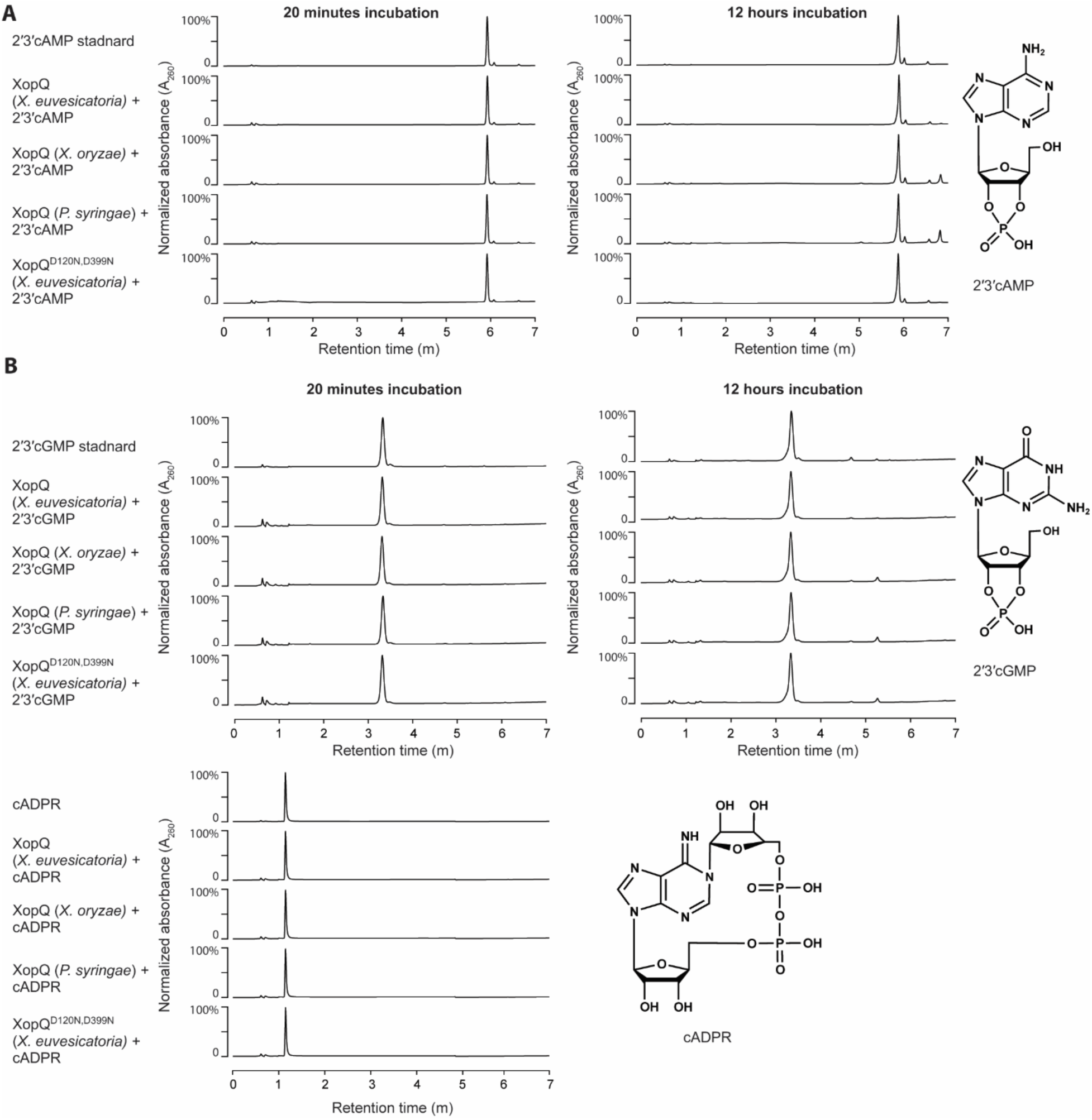
XopQ homologs do not metabolize 2ʹ3ʹcAMP, 2ʹ3ʹcGMP, or canonical cADPR. **A-C**, HPLC chromatograms of in vitro reactions examining the catalytic activity of the indicated XopQ homologs toward 2ʹ3ʹcAMP (**A**), 2ʹ3ʹcGMP (**B**), or canonical cADPR (**C**). In each reaction, 100 nM XopQ and 10 µM molecule were incubated for the indicated duration in room temperature. Data are shown as absorbance values with each chromatogram normalized independently to its maximum value. Chemical structures of the tested compounds are shown. Presented chromatograms are representatives of at least three replicates.

**Figure S6.**
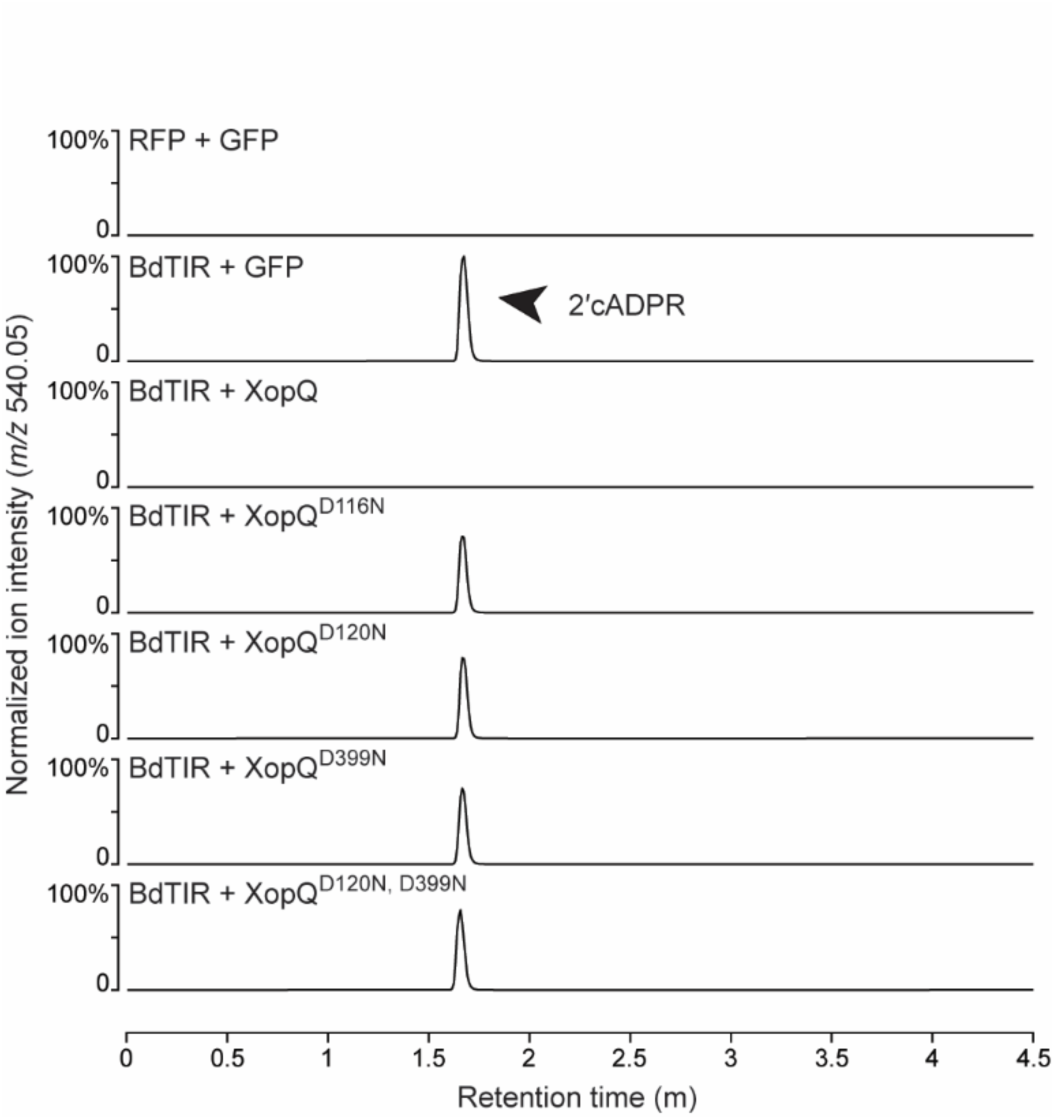
XopQ mutants do not metabolize 2ʹcADPR. HPLC-MS chromatograms of filtered lysates from bacterial strains expressing the indicated proteins. Data are presented as ion intensity for m/z 540.05, with each chromatogram normalized to the maximum value of the BdTIR + GFP control. Shown are representatives of at least three replicates.

**Figure S7.**
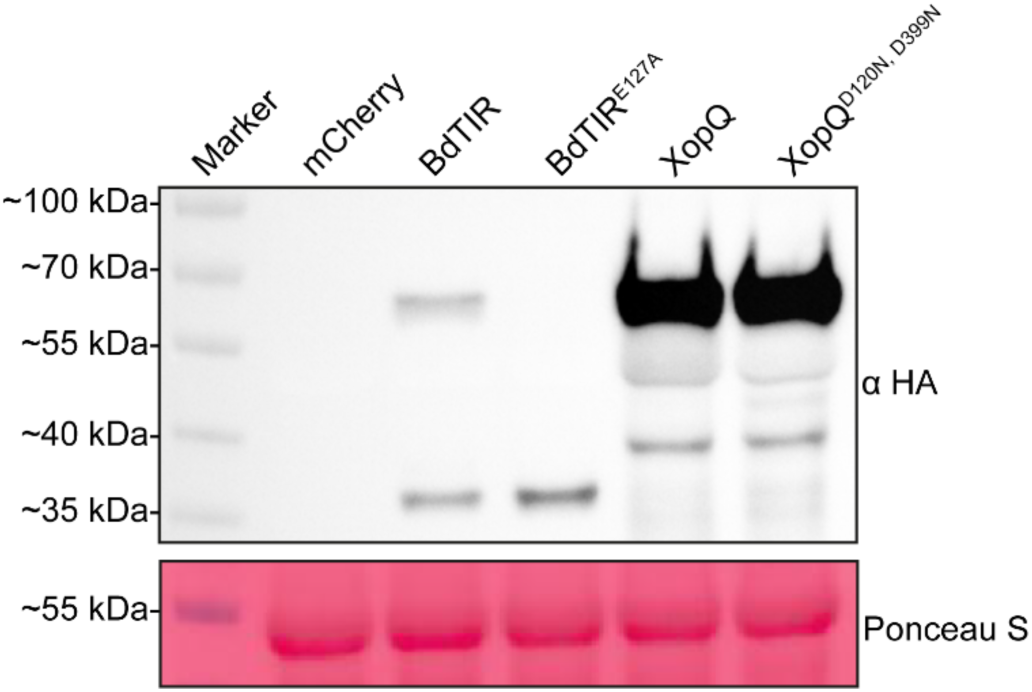
Validation of XopQ and BdTIR overexpression in *N. benthamiana* leaves. Western blot analysis demonstrating expression of the indicated proteins in *N. benthamiana epss* leaves 48 hours post *A. tumefaciens*-mediated transformation. Wild-type and mutant proteins were fused at their N-terminus to triple HA tag, and detection was performed using an anti-HA antibody. Untagged mCherry was used as a negative control.

**Figure S8.**
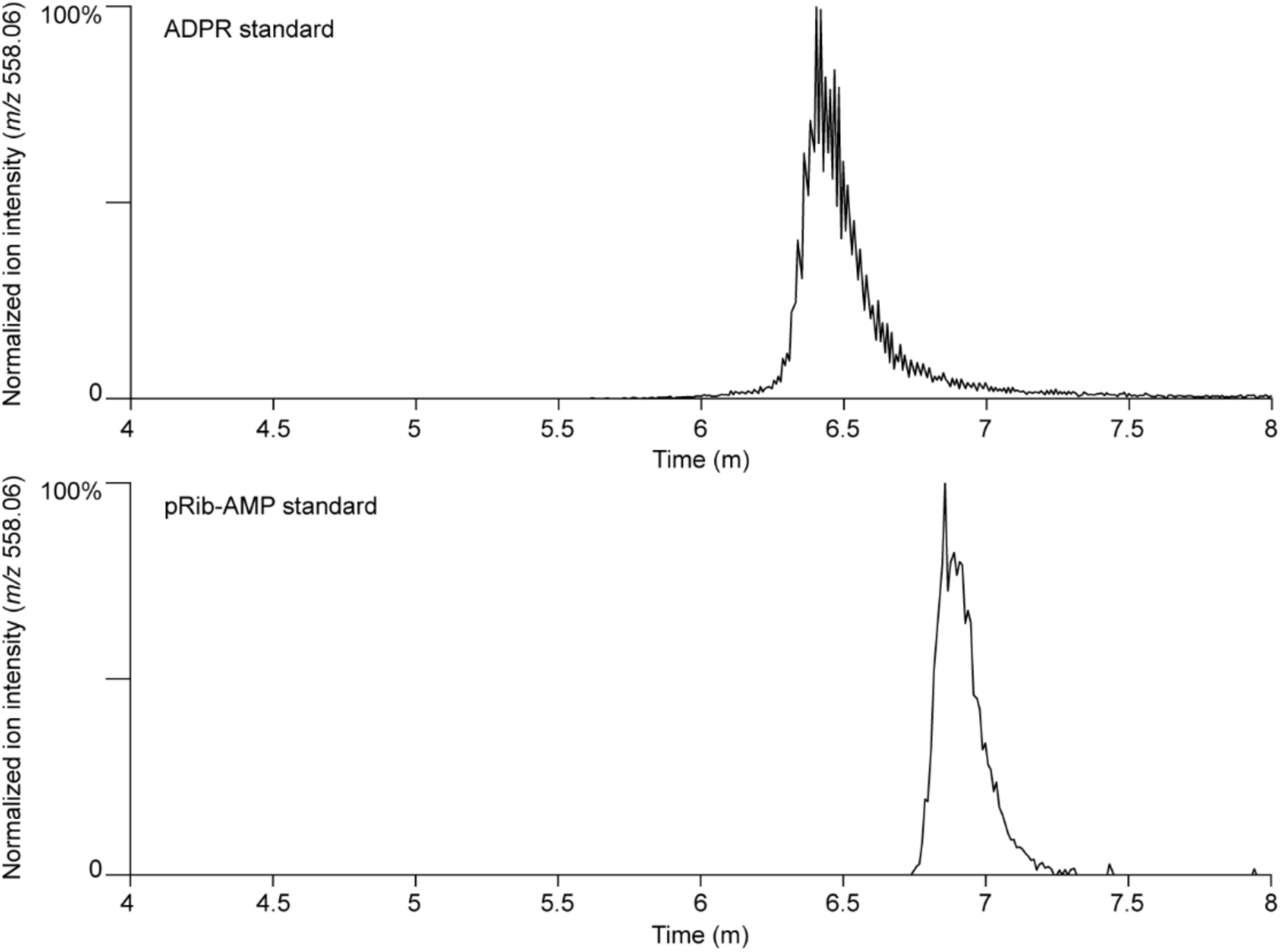
Ion-chromatography mass-spectrometry chromatograms of ADPR and pRib-AMP standards. Data are shown as normalized ion intensities, with each chromatogram normalized independently to its maximum value.

**Figure S9.**
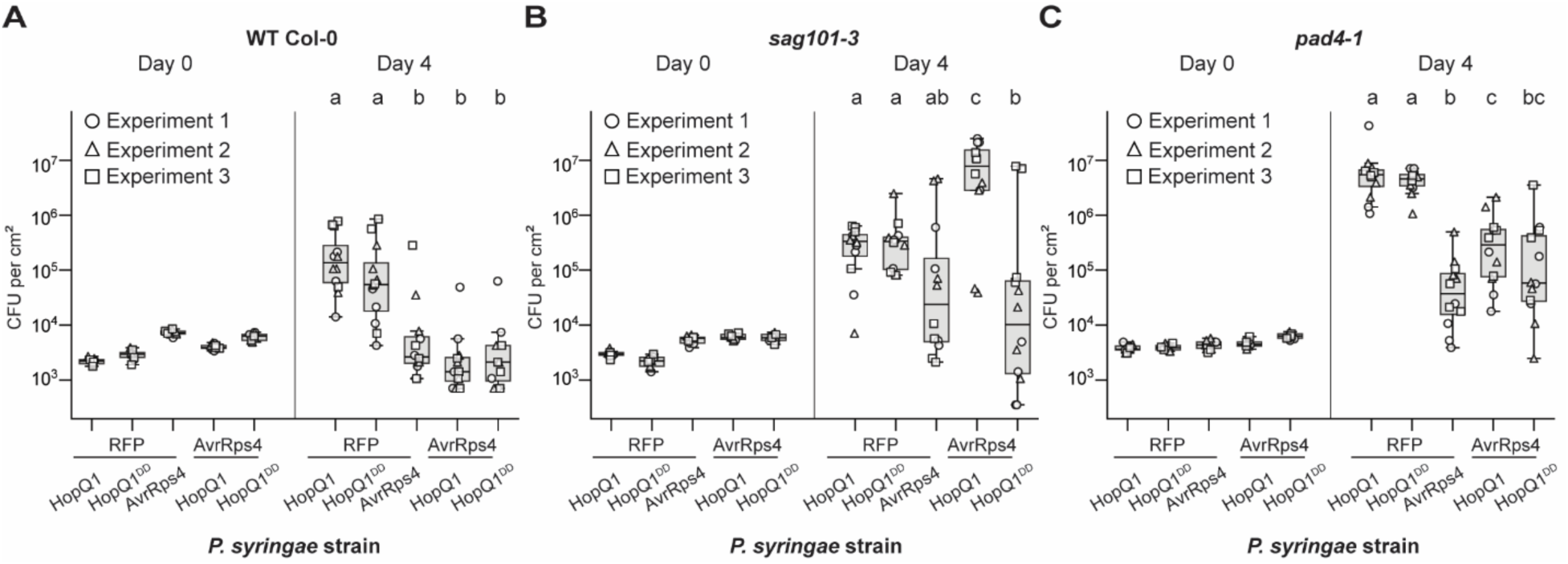
Pathogen-delivered HopQ1 suppresses TIR-mediated PAD4 immunity in *A. thaliana* – complete dataset. Proliferation of *P. syringae* pv. tomato DC3000 Δ*hopQ1* strains carrying operons expressing the indicated proteins in leaves of the *A. thaliana* WT Col-0 (**A**), *sag101-3* (**B**), or *pad4-1* (**C**). HopQ^DD^ denotes the catalytically inactive HopQ^D105N, D384N^. Shown are colony-forming units (CFUs) per cm^2^ measured 1-hour (Day 0) and four days (Day 4) post inoculation from three independent experiments with three (Day 0) or four (Day 4) replicates each (in total, *n* = 9 for Day 0 and *n* = 12 for Day 4). Data are presented as box plots: median (center line), interquartile range (IQR, box), and the most extreme values within 1.5 x IQR (whiskers). Circles, squares and triangles depict individual measurements divided by replicates. Pairwise, two-sided Mann-Whitney U tests followed by Benjamini-Hochberg correction were used to assess differences in bacterial proliferation between strains within each plant genotype. Statistically significant groups, calculated separately for each plant genotype, are indicated via the letters a, b, and c (adjusted P < 0.05). Data in this figure also appears in Figure 5.

